# APLNR regulates directed migration to spatially pattern mesoderm in human gastruloids

**DOI:** 10.64898/2026.10.19.760207

**Authors:** Amanda Brooke Johnson, Shailaja Seetharaman, LiAng Yao, Alex Plum, Zhiyuan Yu, Mattia Serra, Idse Heemskerk, Margaret L. Gardel

## Abstract

GRAPHICAL ABSTRACT

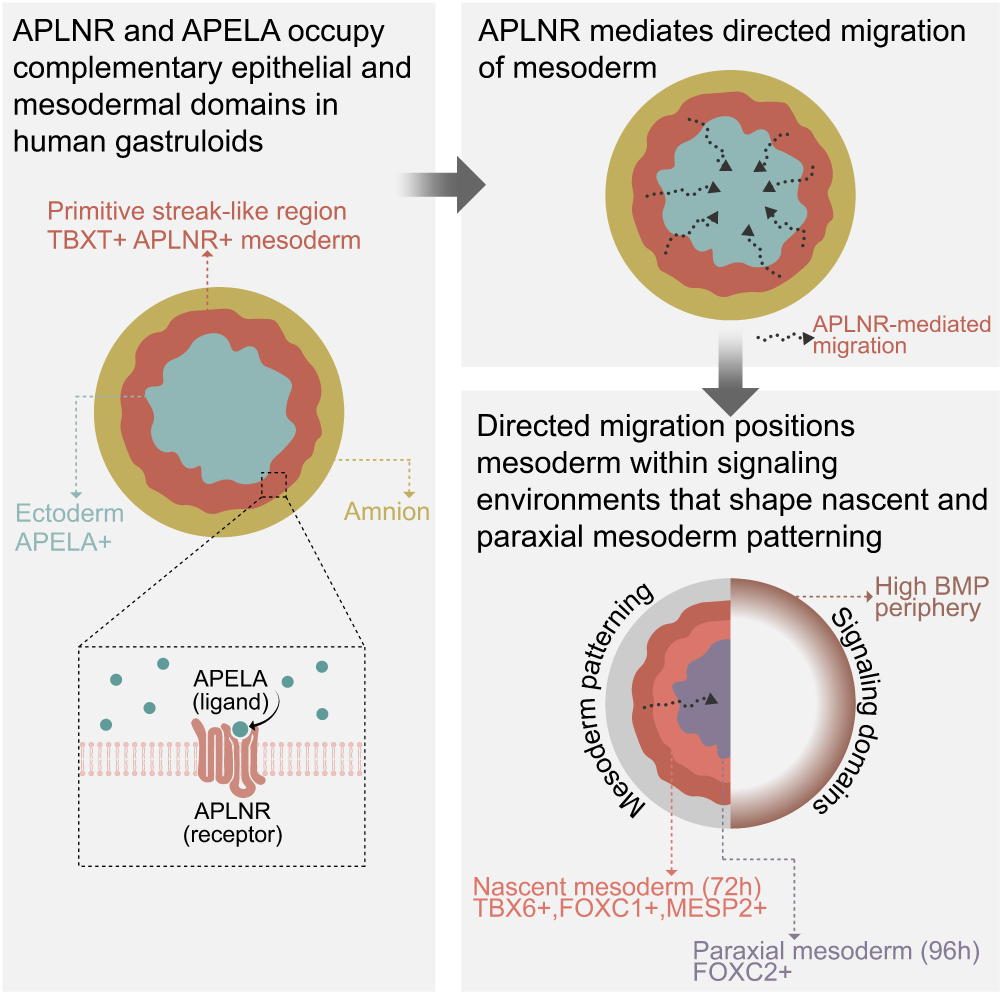

**HIGHLIGHTS:**

- APLNR marks nascent and advanced mesoderm lineages in human gastrulation.
- APELA–APLNR is a candidate paracrine guidance pathway in human gastruloids.
- APLNR is required for directed migration of mesoderm, with little effect on speed.
- Directed migration positions mesoderm for paraxial and lateral differentiation.

During human gastrulation, mesoderm progenitors migrate while acquiring distinct identities. Whether this movement contributes to subsequent patterning remains unclear. Here, we identify the apelin receptor (APLNR) as a regulator of directed migration of mesoderm in human gastruloids. TBXT^+^ cells ingress basally and disperse from the PS-like region toward epiblast-like cells. Tracking reveals that this dispersal reflects a directional bias rather than increased speed. APLNR is enriched in nascent mesoderm, while its ligand APELA is expressed in adjacent epithelial cells. Control gastruloids exhibit fourfold greater dispersal than APLNR-depleted gastruloids despite modest speed differences, consistent with a model of chemotaxis by an epithelial ligand field. Paraxial and lateral mesoderm differentiation competence is preserved following APLNR depletion, while signaling programs and central FOXC2^+^ paraxial organization are altered. These findings support a role for APLNR-dependent guidance in positioning competent mesoderm progenitors for subsequent patterning.

## INTRODUCTION

During mammalian gastrulation, posterior epiblast cells progressively acquire mesendodermal identities as they undergo epithelial-to-mesenchymal transition (EMT) and ingress through the primitive-streak (PS). EMT requires a switch from apical–basal to front–rear polarity to enable migration beneath the epiblast and disperse away from the PS^1–5^. Mesoderm migration involves both the acquisition of cell motility and the regulation of migration direction^6,7^. Motogenic signals can promote cell movement without specifying direction^8^, whereas guidance cues bias migration toward specific locations^7^. Whether these processes are independently regulated during human gastrulation remains unclear.

Directed mesoderm migration during gastrulation can be regulated by chemical guidance cues. In chick embryos, fibroblast growth factor 8 (FGF8) produced near the PS repels ingressing mesoderm, whereas FGF4 from the forming notochord attracts mesoderm toward the midline^7^. In mouse embryos, cells lacking *Fgf8* undergo EMT but largely fail to migrate away from the PS^9^. Chemotactic guidance during gastrulation can also be mediated by G protein-coupled receptors (GPCRs). In zebrafish, the apelin receptor APLNR and its ligand APELA (also known as ELABELA or Toddler) regulate mesendoderm movement towards the animal pole^10,11^. Although APELA is initially produced throughout the blastoderm, migrating APLNR-positive cells locally deplete the ligand, producing a self-generated gradient that guides their movement^12^. In human stem cell models, APELA is secreted by pluripotent cells, whereas APLNR is induced during mesendoderm differentiation^13^. APLNR has also been implicated in mesoderm development and cell outgrowth in human embryoid bodies^14^. However, whether APLNR-dependent guidance operates during human gastrulation remains unknown.

Directed mesoderm migration is thought to contribute to fate specification and tissue patterning during gastrulation^15–17^. However, separating the contributions of migration and fate specification to mesoderm patterning in intact embryos is challenging because regulators of migration often affect other cellular processes^16,18^. In mouse embryos, disruption of the small GTPase Rac1 or its downstream WAVE regulatory complex causes migration and mesoderm patterning defects^18,19^. However, Rac1 also regulates adhesion, survival, and intracellular signaling, so these phenotypes cannot be attributed to migration alone^18^. Likewise, loss of APELA in mice affects early APLNR-positive mesodermal derivatives and causes cardiovascular defects that could reflect altered migration or direct effects on developmental signaling^20^. These overlapping functions motivate simpler systems in which migration can be measured alongside differentiation and tissue patterning.

Micropatterned two-dimensional (2D) human gastruloids recapitulate the spatial organization of peri-gastrulation embryos, forming a TBXT (Brachyury)-positive (TBXT^+^) domain adjacent to SOX2^+^ epiblast-like cells^21–23^. Their defined geometry and reproducible patterning provide a tractable system for investigating the relationship between mesoderm migration, signaling, and differentiation^24–28^. Previous studies have used 2D gastruloids to map cell migration and fate^29^. Extending culture to 96 hours revealed that inward migrating mesoderm cells contribute to central paraxial-like domains, while FOXF1^+^ lateral mesoderm forms near the colony edge where BMP signaling is highest^30^. The mechanisms that direct TBXT^+^ mesoderm migration and the contribution of this movement to subsequent mesoderm patterning remain unresolved.

Here, we employ human stem cell-derived gastruloids to investigate the mechanisms directing mesoderm migration and how this movement contributes to subsequent fate specification and tissue patterning. Tracking TBXT^+^ cells as they disperse from the PS-like region toward epiblast-like cells reveals a persistent directional bias that is largely separable from cell speed. Human embryo transcriptomes and gastruloid data reveal APLNR expression in nascent mesoderm alongside APELA expression in neighboring epithelial compartments. APLNR depletion disrupts directed migration, while a minimal chemotaxis model suggests that an epithelial APELA source could account for the observed dynamics. Under directed differentiation conditions, APLNR-depleted progenitors retain broad competence to form paraxial and lateral mesoderm. In gastruloids, however, they exhibit altered signaling and mesoderm patterning programs together with reduced central organization of FOXC2^+^ paraxial mesoderm. These findings support a model in which APLNR-dependent directed migration positions mesoderm progenitors within signaling environments that shape subsequent patterning.

## RESULTS

### Human gastruloid mesoderm undergoes basal ingression and directionally biased dispersal

Human pluripotent stem cells confined to circular micropatterns 500–1000 *µ*m in diameter reproducibly form radially organized germ-layer domains following treatment with bone morphogenetic protein 4 (BMP4)^21^. To generate gastruloids, cells were plated as epithelial monolayers on circular micropatterns and treated with 50 ng/mL BMP4 at *t* = 0 in mTeSR^21^. All exogenous cues are supplied uniformly, so the resulting spatial organization arises from the cells rather than from an externally applied gradient. By 48 h, gastruloids are organized into concentric domains: ISL1^+^ amnion-like cells at the outer edge, a dense TBXT^+^ primitive-streak-like (PS-like) annulus just inside them, and SOX2^+^ epiblast-like cells at the center. We distinguish two phases within the first 48 h of differentiation: establishment of the PS-like region between 24 and 36 h, followed by mesoderm ingression and dispersal between 36 and 48 h (Figure 1A).

**Figure 1.**
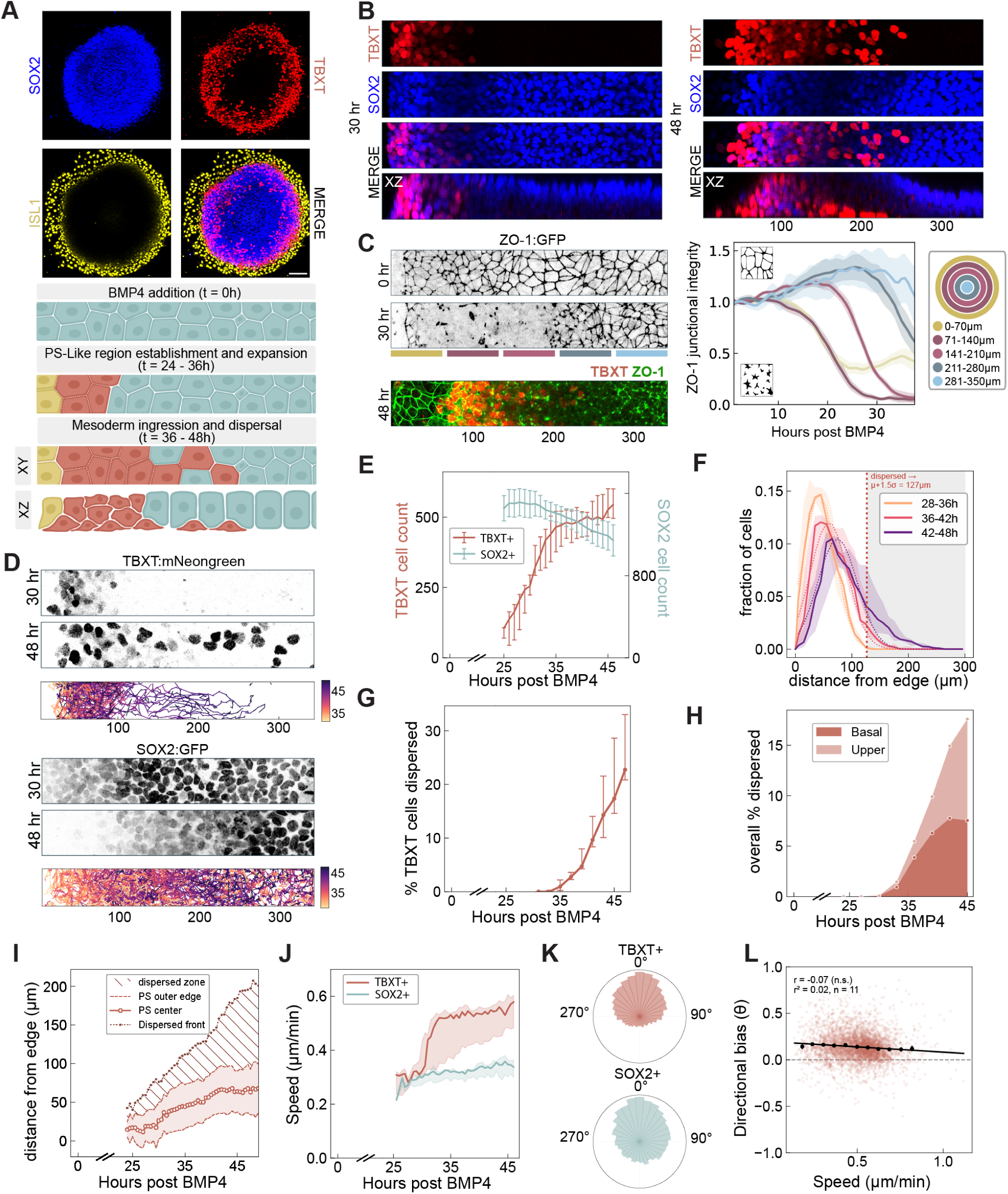
Human mesoderm undergoes basal ingression and directed dispersal from a PS-like region. **(A)** Representative 48-h gastruloid stained for TBXT, SOX2, ISL1, and DAPI, with schematic of the micropattern assay after BMP4 (*t* = 0); XY and XZ views show the radial and epithelial-depth axes. Scale bar, 100 *µ*m. **(B)** Representative top-view (xy) and orthogonal (xz) sections at 30 and 48 h post-BMP4, stained for TBXT, SOX2, and DAPI; numbers, distance from the colony edge. **(C)** Radial strips of live ZO-1–GFP gastruloids at 0, 24, and 36 h (left) and ZO-1 remodeling front position over time (right); 48-h ZO-1/TBXT inset from Figure S1A. *n* = 6, *N* = 2 (Video S1). **(D)** Live-imaging stills and time-colored single-cell trajectories of TBXT^+^ and SOX2^+^ reporter gastruloids (Videos S2, S3). Track duration distributions are shown in Figure S1F. **(E)** Tracked TBXT^+^ and SOX2^+^ cells per imaging frame. Median and interquartile range (IQR) across gastruloids; points, individual gastruloids. TBXT^+^: *n* = 10, *N* = 3; SOX2^+^: *n* = 4, *N* = 2. **(F)** Radial density of TBXT^+^ cells over time, with Gaussian fit at the final time point. PS-like boundary, *µ* + 1.5*σ* (127 *µ*m from the colony edge; Figure S1B). *n* = 10, *N* = 3. **(G)** Fraction of TBXT^+^ cells dispersed beyond the PS-like boundary in three time windows. Points, individual gastruloids; bars, median. *n* = 10, *N* = 3. **(H)** Dispersed TBXT^+^ cells in basal and upper z-planes over time, relative to all TBXT^+^ cells. *n* = 4, *N* = 1 (Figure S1C; Video S4). **(I)** PS-like region over time (solid line, Gaussian center *µ*; shading, *µ ± σ*; dashed line, boundary *µ* + 1.5*σ*) and dispersed front (90th percentile of cells beyond the boundary). *n* = 10, *N* = 3. **(J)** Migration speed of TBXT^+^ and SOX2^+^ cells by time window. Points, individual gastruloids; bars, median; Mann–Whitney tests (Methods). TBXT^+^: *n* = 10, *N* = 3; SOX2^+^: *n* = 4, *N* = 2. **(K)** Rose plots of net start-to-end displacement angles (0° inward, 180° outward) for trajectories with at least five edges. TBXT^+^: 17,927 tracks, *n* = 10, *N* = 3; SOX2^+^: 12,826 tracks, *n* = 4, *N* = 2. Per-gastruloid mean resultant lengths in Figure S1E. **(L)** Speed versus inward orientation of TBXT^+^ cells with at least 10 tracked edges; per-gastruloid Pearson correlation, one-sample Wilcoxon signed-rank test (Methods). 733 cells, *n* = 10, *N* = 3.

TBXT marks both PS-like and nascent mesoderm states. By 48 h, a subset of TBXT^+^ cells are located beyond the dense PS-like annulus toward the colony center. Orthogonal sections at 30 and 48 h show TBXT^+^ cells beneath the SOX2^+^ epiblast-like layer by 48 h, which is consistent with mesoderm ingression (Figure 1B).

To examine epithelial remodeling during early gastruloid differentiation, we performed live imaging of the tight-junction protein ZO-1 using a ZO-1–GFP reporter^31^. We quantify junctional organization from skeletonized ZO-1 networks within consecutive 70 *µ*m radial bins from the colony edge (Figure 1C; Video S1). ZO-1 initially forms a continuous junctional network that progressively remodels inward, reaching approximately 200 *µ*m from the colony edge by 36– 38 h. By 48 h, ZO-1 is reduced throughout the TBXT^+^ domain but remains prominent in the SOX2^+^ epithelial compartment and ISL1^+^ amnion-like domain (Figure S1A). This is consistent with the epithelial and mesenchymal marker organization previously described in micropatterned gastruloids, including E-cadherin, N-cadherin, EPCAM and SNAI1^30,32^. Epithelial remodeling therefore begins during establishment of the PS-like region.

To resolve mesoderm dynamics relative to the epiblast-like compartment, we imaged TBXT– mNeonGreen^33^ and SOX2–GFP^31^ reporter gastruloids from 24 h, during establishment of the PS-like region (Figure 1D; Videos S2, S3). TBXT^+^ cells appear by 25 h and rapidly increase in number until 35–36 h, after which the population largely plateaus. SOX2^+^ cell number remains initially stable and then declines by approximately 25–30% between 34 and 46 h (Figure 1E). TBXT^+^ cells initially form a dense annulus with few cells toward the colony center. From approximately 36 h, cells begin leaving this PS-like region and move towards the epiblast region (Figure 1D,F). We refer to this movement as mesoderm dispersal and quantify it using a fixed inward boundary derived from the TBXT^+^ radial density profile (Methods). Dispersal increases from approximately 35 to 46 h, with a median of 20.6% of TBXT^+^ cells lying beyond the PS-like region during 42–48 h (Figure 1G).

To quantify the apical–basal distribution of TBXT^+^ cells, we assign cells to basal or upper layers according to nuclear center Z-position in stacks of 11 planes spaced 3.5 µm apart. The first plane was positioned at the nuclei nearest the glass substrate and defines the basal layer; the remaining planes define the projected upper layers. Tissue organization changes from an initially pseudostratified epithelium to approximately four to five cell layers by 48 h (Figure 1B). Because Z-sampling is limited during long-term live imaging, we track cells independently within each layer(s). Dispersed TBXT^+^ cells accumulate predominantly in the basal layer, with a smaller contribution from the upper layers (Figure 1H; Video S4). This enrichment is not explained by unequal TBXT^+^ cell abundance across layers (Figure S1C). Together with the orthogonal sections, these data indicate that mesoderm ingresses beneath the epiblast and disperses inward primarily within the basal layer. Radial density profiles further reveal two concurrent changes: the dense PS-like region shifts progressively inward while maintaining a similar width, and increasing numbers of TBXT^+^ cells disperse beyond this region toward the gastruloid center (Figure 1I; Video S4). These measurements distinguish movement of the PS-like region itself from mesoderm dispersal beyond it.

TBXT^+^ cells accelerate as dispersal begins, with speed increasing approximately twofold from *∼* 0.3 *µ*m/min at 25–34 h to *∼* 0.6 *µ*m/min at 35–48 h. At later times they move faster than SOX2^+^ epiblast-like cells (Figure 1J). Yet dispersed TBXT^+^ cells do not move significantly faster than those remaining within the PS-like region (Figure S1D). Their dispersal instead accompanies preferentially inward migration: TBXT^+^ trajectories are more strongly oriented toward the center than SOX2^+^ trajectories and have 45% higher mean resultant length across gastruloids (Figure 1K; Figure S1E). TBXT^+^ cells show net inward displacement (Figure S1G), but faster tracks are not more inwardly oriented within individual gastruloids (Figure 1L). Mesoderm dispersal therefore reflects an inward directional bias that is separable from cell speed.

### APELA and APLNR occupy complementary epithelial and mesodermal compartments

During human gastrulation mesoderm emerges from the PS and spreads anteriorly and laterally between the epiblast and hypoblast. As development proceeds, the remaining epiblast forms ectoderm, mesoderm establishes axial, paraxial, and lateral territories, and definitive endoderm replaces the hypoblast (Figure 2A). Carnegie Stage 7 (CS7) and Carnegie Stage 8 (CS8) capture successive phases of this process, from mesoderm production within the PS to its expansion between the ectodermal and endodermal layers^34^.

**Figure 2.**
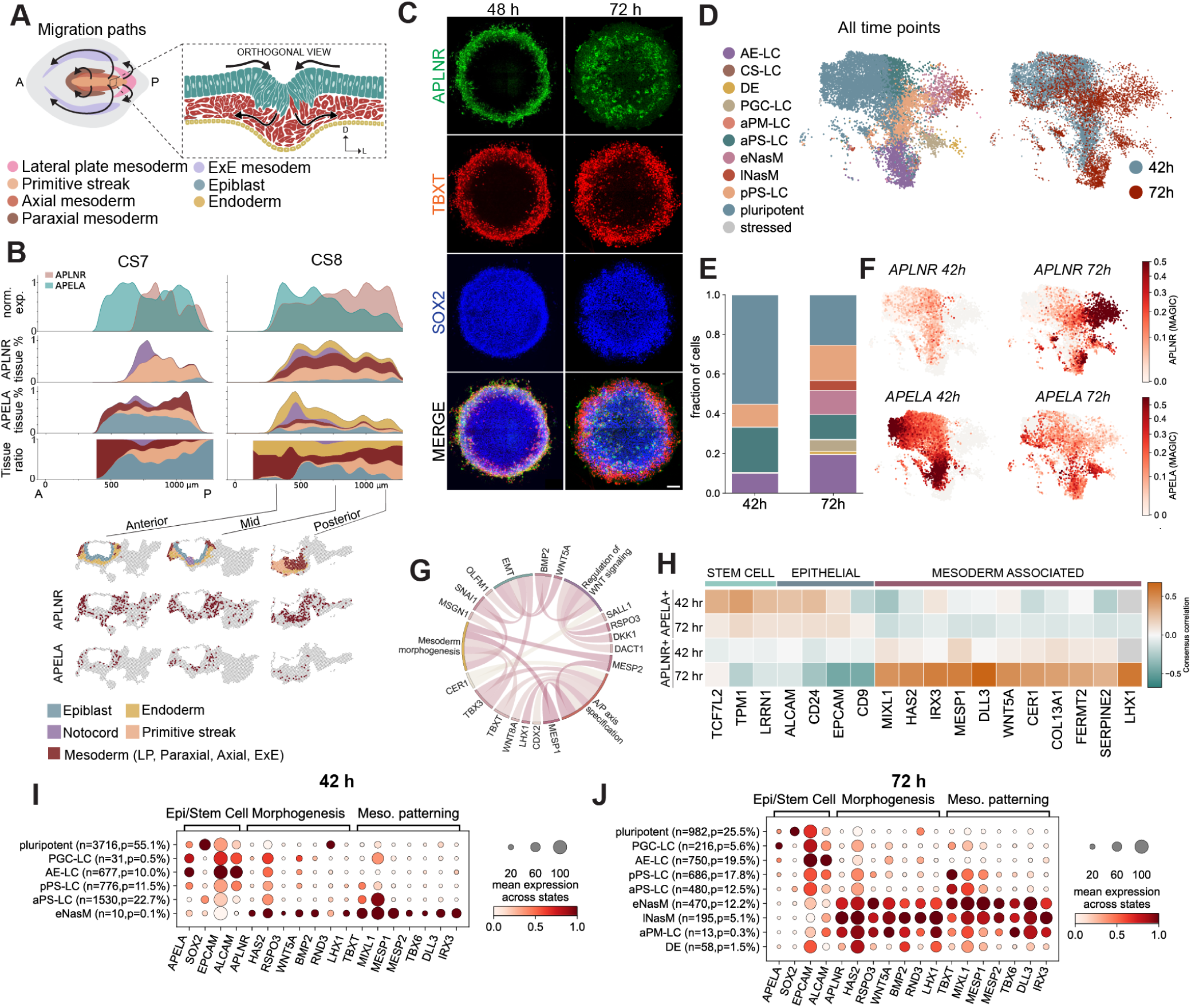
APELA and APLNR occupy complementary epithelial and mesodermal compartments. **(A)** Schematic of mesoderm migration during gastrulation: emergence through the PS and dispersal into axial, paraxial, lateral plate, and extraembryonic territories, with orthogonal view of epiblast, mesoderm, and endoderm. **(B)** Published Stereo-seq data from human Carnegie stage 7 (CS7) and 8 (CS8) embryos. Top, normalized *APLNR* and *APELA* expression across serial anterior–posterior slices. Bottom, expression in representative anterior, mid, and posterior sections; tissue annotation by color. CS7 *APELA* was imputed from the matched single-cell reference; all other profiles are observed. Profiles are smoothed along the anterior–posterior axis (*σ* = 2 sections) and scaled to a maximum of 1 per gene and stage; magnitudes are not comparable between genes or stages. **(C)** Whole-mount immunofluorescence of human gastruloids at 48 and 72 h stained for APLNR, TBXT, and SOX2. Scale bar, 100 *µ*m. **(D)** UMAP of control gastruloid single-cell RNA-seq at 42 and 72 h, colored by cell state (left) or time point (right). *N* = 2. **(E)** Relative abundance of cell states at 42 and 72 h. *N* = 2; per-repeat counts and proportions in Table S2. **(F)** UMAP feature plots of *APLNR* and *APELA* at 42 and 72 h. *N* = 2. **(G)** Gene Ontology biological processes enriched in the 218-gene APLNR-containing co-expression module (WGCNA, control mesoderm cells); ribbons connect module genes to significantly enriched processes. *N* = 2. Module membership (*kME*) and hub-gene rank in Table S3. **(H)** Correlation of *APELA* and *APLNR* with representative stem-cell, epithelial, and mesoderm-associated genes at 42 and 72 h; color, consensus correlation across biological repeats (Methods). *N* = 2. **(I,J)** Dot plots of *APELA*, epithelial markers, *APLNR*, and patterning and morphogenesis genes from the APLNR-containing module across cell states at 42 h (I) and 72 h (J). Dot size, fraction of cells with detectable expression; color, mean expression scaled 0 to 1 per gene within each time point. *N* = 2; cell numbers per state in row labels.

Several ligand–receptor pathways could plausibly guide mesoderm in the human embryo. We apply LIANA rank aggregation to published single-cell RNA-sequencing data from CS7^35^ and CS8^36^, focusing on predicted paracrine interactions from other cell populations to mesoderm. Among signaling families with known roles in migration, EMT, and guidance, APELA*→*APLNR is recovered at both stages, with endoderm and epiblast/ectoderm as the predominant sending populations (Table S1). At CS8, *APELA* was detected in only 5.8% of mesoderm cells and is more prevalent in epithelial compartments. APELA–APLNR signaling therefore emerges as a plausible guidance pathway, consistent with its established role in mesoderm migration in other vertebrates^12,37^.

Published Stereo-seq data reveal complementary spatial distributions of *APLNR* and *APELA* in CS7 and CS8 human embryos^36,38^ (Figure 2B). At CS7, *APLNR* expression is concentrated posteriorly in the PS. Because *APELA* was absent from the retained spatial gene set, we imputed its distribution using a CS7 single-cell RNA-sequencing reference^35^. Imputed *APELA* expression is concentrated in the epiblast/ectodermal compartment and peaks anterior to *APLNR*. By CS8, *APLNR* expression extends beyond the PS into mesodermal territories, while directly measured *APELA* expression remains concentrated in endodermal and epiblast/ectodermal compartments.

In gastruloids APLNR protein is enriched in TBXT^+^ cells relative to the SOX2^+^ epithelium. It is concentrated in the PS-like region at 48 h and present in dispersed TBXT^+^ cells by 72 h (Figure 2C). Single-cell RNA sequencing at 42 and 72 h resolves pluripotent, epithelial-associated, PS-like, and nascent mesoderm populations (Figure 2D,E). We annotate anterior and posterior PS-like cells (aPS-LC and pPS-LC) and early and late nascent mesoderm (eNasM and lNasM) using a published gastruloid reference^30^. Canonical markers support these annotations across biological repeats (Figure S2A–E; Table S7). At 42 h, *APELA* is concentrated in pluripotent and epithelial-associated populations while *APLNR* is enriched in PS-like and emerging mesodermal populations. By 72 h, *APLNR* is concentrated in nascent mesoderm while *APELA* remains primarily epithelial (Figure 2F). This mirrors the compartmental organization seen in the CS7 and CS8 human embryo datasets.

Weighted gene co-expression network analysis places *APLNR* within a module enriched for mesoderm morphogenesis, anterior–posterior patterning, EMT, and WNT regulation (Figure 2G; Figure S2F–H). Highly connected genes include the mesoderm regulators *MESP1* and *TBX6* alongside signaling and cell-behavior genes such as *WNT5A*, *HAS2*, and *RND3*. APLNR expression therefore accompanies a program linking mesoderm development with the machinery for tissue reorganization. Gene-wise correlations support this association: *APELA* correlates with stem-cell and epithelial genes at 42 h, whereas *APLNR* correlates positively with mesoderm patterning and morphogenesis genes and negatively with epithelial genes at 72 h (Figure 2H).

The state-resolved expression profiles reveal a further distinction between PS-like cells and nascent mesoderm (Figure 2I,J). At 72 h, *APLNR* is detected in approximately 83% of eNasM and 90% of lNasM cells compared with only 19% of aPS-LC and 18% of pPS-LC cells. APLNR expression is thus most prevalent after cells acquire nascent mesoderm identities, while *APELA* remains enriched in epithelial-associated populations. This separation of ligand and receptor expression is consistent with epithelial APELA providing a guidance cue for emerging mesoderm.

### APLNR regulates directed mesoderm migration

The complementary spatial organization of APLNR and APELA identified APLNR as a candidate regulator of mesoderm guidance. To test whether APLNR is required for directed migration, we generated a doxycycline-inducible^39,40^ shAPLNR line in the TBXT–mNeonGreen hESC cell line^33^. APLNR knockdown reduces transcript abundance by a median of 80% (Figure 3A) and shows lower APLNR localization within TBXT^+^ cells (Figure 3B). The relative organization of the TBXT^+^ and SOX2^+^ compartments is preserved and the peripheral ISL1^+^ amnion domain remains present (Figure S3E). Live imaging shows that shAPLNR TBXT^+^ cells remain concentrated within the PS-like region by 48 h post-BMP4 (Figure 3C; Video S5). TBXT^+^ cell number is similar between conditions (Figure 3D), indicating that the different cell distributions do not reflect fewer TBXT^+^ cells.

**Figure 3.**
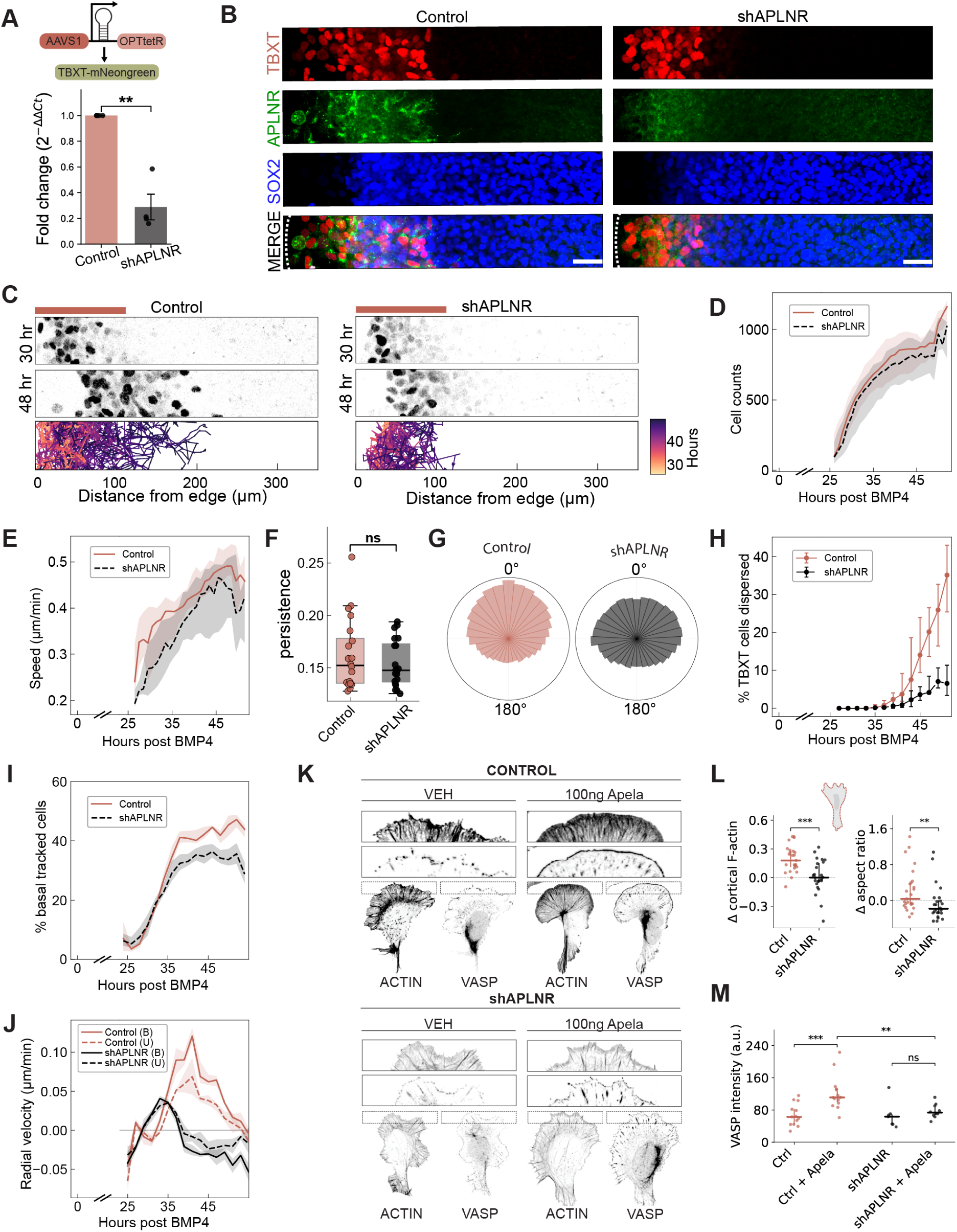
APLNR regulates directed mesoderm migration. **(A)** shAPLNR construct and APLNR knockdown validation by qPCR; one-sample *t*-test, *N* = 4. **(B)** Representative immunofluorescence of control and shAPLNR gastruloids at 48 h post-BMP4 stained for TBXT, APLNR, and SOX2. Scale bar, 100 *µ*m. **(C)** Live-imaging stills and time-colored TBXT^+^ trajectories of control and shAPLNR gastruloids; pink bars, PS-like region (Video S5). Representative of *n* = 20 per condition, *N* = 4. **(D)** Tracked TBXT^+^ cells per gastruloid over time; paired *t*-test at the replicate level. *n* = 20 per condition, *N* = 4. **(E)** Migration speed of TBXT^+^ cells over time (per-cell mean track speed); median across gastruloids in 1-h bins, shading IQR. Replicate level in Figure S3H. *n* = 20 per condition, *N* = 4. **(F)** Persistence of TBXT^+^ trajectories; points, individual gastruloids; paired Wilcoxon signed-rank test at the replicate level (Figure S3I). *n* = 20 per condition, *N* = 4. **(G)** Rose plots of net displacement angles (as in Figure 1K) for trajectories with at least five edges: 62,476 control and 57,705 shAPLNR tracks; *n* = 20 per condition, *N* = 4. Replicate-level mean resultant lengths in Figure S3G. **(H)** Fraction of TBXT^+^ cells dispersed beyond the PS-like boundary (*µ* + 1.5*σ* of the control fit; 112 *µ*m from the colony edge) over time; mixed-effects model (Methods). *n* = 20 per condition, *N* = 4. **(I)** Fraction of tracked mesoderm cells in the basal compartment over time; median and IQR across gastruloids. *n* = 4 per condition, *N* = 1. **(J)** Radial velocity of basal (solid) and upper (dashed) mesoderm over time; positive values, inward. Median and IQR across gastruloids. *n* = 4 per condition, *N* = 1. **(K)** Representative F-actin and VASP staining of control and shAPLNR mesoderm progenitors treated with vehicle or 100 ng/mL ELA-32 for 5 min. **(L)** Cortical F-actin enrichment and aspect ratio of control and shAPLNR progenitors after APELA, relative to vehicle. Points, individual APELA-treated cells; lines, median and IQR. *N* = 3; *n* cells (vehicle, APELA): control 27, 25; shAPLNR 19, 24. **(M)** Peripheral VASP intensity in control and shAPLNR progenitors treated with vehicle or 100 ng/mL ELA-32. Points, individual cells; lines, median and IQR. *N* = 1, *n* = 44 cells.

shAPLNR cells remain motile and follow paths of similar straightness to controls, although their speed is modestly lower (Figure 3E,F; Figure S3H). Control TBXT^+^ trajectories are preferentially oriented toward the gastruloid center, whereas shAPLNR trajectories show weaker inward orientation and more tangential movement (Figure 3G; Figure S3D). Mean resultant length is lower in shAPLNR gastruloids across all four biological repeats (Figure S3G). APLNR thus supports the inward orientation of migration without substantially changing trajectory straightness.

Control gastruloids show approximately fourfold greater mesoderm dispersal than shAPLNR gastruloids: 20.3% of tracked TBXT^+^ cells cross beyond the PS-like boundary between 42 and 48 h compared with 5.1% following APLNR depletion (Figure 3H). This difference remains significant after accounting for migration speed, supporting a contribution of APLNR to dispersal beyond its modest effect on speed (Figure 3H; Methods). The effect is consistent across all four biological replicates and persists when the boundary threshold is varied from 1*σ* to 2*σ* (Figure S3A,B,F). The dense PS-like region also moves farther inward in controls (Figure S3C). APLNR therefore supports both inward translocation of the PS-like region and mesoderm dispersal beyond it.

APLNR depletion separates basal occupancy from inward migration. Basal cells remain a substantial fraction of tracked mesoderm in both conditions between 42 and 48 h (Figure 3I; Videos S6, S7). In controls, basal cells migrate inward with 1.6-fold greater radial velocity than upper cells. Inward migration is reduced in both compartments following APLNR depletion (Figure 3J; Videos S6, S7). The dispersal defect therefore persists even among cells positioned beneath the epiblast-like layer.

We then examined whether APELA engages the protrusive machinery of mesoderm progenitors. Actin assembly at the cell periphery supports lamellipodia that propel and steer migrating cells^41^. In TBXT^+^ progenitors generated by early mesoderm induction, acute ELA-32 stimulation increases peripheral F-actin enrichment in controls but produces little change in shAPLNR cells (Figure 3K,L). The actin-assembly regulator VASP also accumulates at the cell periphery in stimulated controls, with no detectable increase following APLNR depletion (Figure 3K,M). These responses connect APELA–APLNR signaling to the cytoskeletal machinery used for cell migration.

### A minimal chemotaxis model reproduces APLNR-dependent mesoderm dispersal

A spatially varying APLNR ligand field could in principle account for the observed PS dispersal (Figure 4A). Because APELA was the predominant APLNR ligand transcript in gastruloids (Figure S2G) and was enriched in epithelial populations, we modeled APELA as a candidate spatial cue. We considered a radially symmetric continuum model in which mesoderm exhibits directed motility with velocity **v** and undirected motility with effective diffusivity *D_m_*. To capture upper mesoderm’s lower inward radial velocity (Figure 3J), we modeled separate upper and basal areal densities *m_u_*(*r, t*) and *m_b_*(*r, t*) and velocities **v***_u_* = *α***v***_b_* (*α* = 0.5), and ingression from upper to basal at rate *γ*:

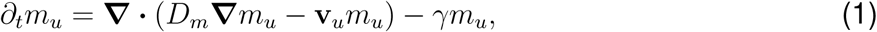

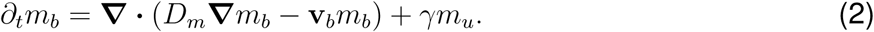

**Figure 4.**
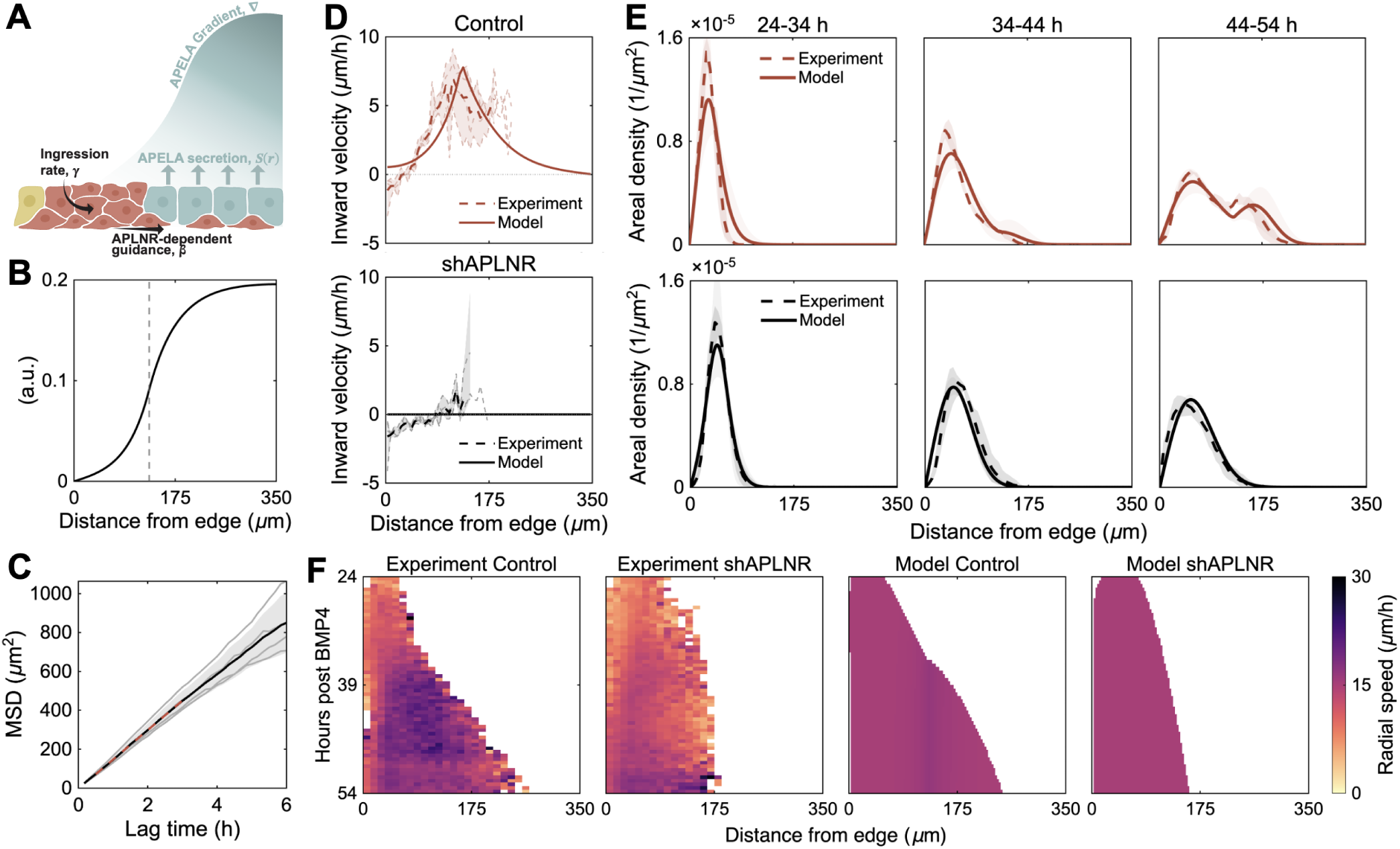
Minimal chemotaxis model reproduces mesoderm dispersal. **(A)** Schematic of the control condition. APELA produced by a central SOX2^+^ source establishes a radial concentration gradient (blue). Mesoderm exhibits random motility and directed migration up the gradient, which is stronger for basally ingressed cells (dark red). **(B)** Steady-state APELA concentration profile. Dashed line marks the source boundary *R − r_s_*. **(C)** Mean-squared displacement (MSD) versus time lag across APLNR knockdown replicates (gray lines). Dashed red line shows a linear fit to the mean MSD over 0.5 *−* 3 h. Replicate specific fits give an effective mesoderm diffusivity *D_m_* = 37.9 *±* 4.6*µ*m^2^*/*h (mean*±*SD, *n* = 4). **(D-F)** Model-experiment comparisons for control (red) and knockdown (black). **(D)** Experimental and modeled inward radial-velocity versus radial position, averaged over 24–54 h (Methods). Dashed lines show the replicate mean and shading individual experimental replicates and the range. Red lines show density-weighted model velocity averaged over 24 *−* 54 h. Experimental curves reflect *n* = 4. See Figure S4E for inward velocities over each 10 h interval. **(E)** Experimental and model TBXT+ densities for control (*n* = 4) and knockdown (*n* = 4). Experimental curves (dashed) and bands show the mean and range across replicates after averaging within each time interval. Model curves (solid) and bands show the mean and range within each time interval. **(F)** Experimental and model heatmaps of radial speed over time for TBXT+ cells. Experimental heatmaps were calculated from tracked cell displacements and averaged across replicates. Model heatmaps show expected radial speed from directed and undirected motility.

Basal directed motility **v***_b_* = *β****∇****c* follows the gradient of APELA concentration *c*(*r, t*).

APELA concentrations could not be measured directly, so we modeled APELA as produced by a SOX2^+^ source *S*(*r*) with uniform diffusion *D_c_*and clearance *k*. Estimates of *D_c_* and *k* for other secreted signaling molecules in 2D gastruloids^24^ provided order-of-magn<u>itude</u> references. With *D_c_*= 10^4^ *µ*m^2^*/*h and *k* = 5 h*^−^*^1^, the characteristic gradient length 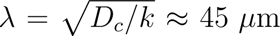 is short relative to the colony size, and turnover is fast relative to the 30-h modeling interval, so we considered a steady-state profile satisfying *S*(*r*) + *D_c_****∇***^2^*c − kc* = 0 (Figure 4B). The gradient is further shaped by *S*(*r*) (Figure S4A) and the outer boundary condition. We modeled *S*(*r*) as a step function from 0 to 1 at a source boundary placed 130 *µ*m from the colony edge (dashed line in Figure 4B), approximately circumscribing the region of APELA-expressing SOX2^+^ cells and lying near the PS-like boundary. We modeled an absorbing outer colony boundary for APELA (*c* = 0), and a no-flux condition yields a similar gradient (Figure S4D).

Because both upper and basal cells in shAPLNR gastruloids lose most of their net inward velocity (Figure 3J), we used their mean-squared displacement to upper-bound the effective diffusivity *D_m_* (Figure 4C), conservatively setting *D_m_*= 30*µ*m^2^*/*h in case of residual directed motion. We modeled APLNR knockdown by setting *β* = 0, leaving only undirected motility. In controls, a range of *D_m_* suffices to reproduce density profiles, provided sufficiently high *β* (Figure S4B). To constrain *β*, we examined radial-velocity profiles (Figure 4D). The linear response **v** *∝ **∇**c* predicts a peak inward velocity where the ligand gradient is steepest, near the source boundary. This peak was present in the time-averaged radial-velocity profiles of all four control replicates and its magnitude (Figure 4D) provided a reference for *β*.

Starting from initial conditions fit to average early areal densities (Figure S4F), the model reproduces the evolution of mesoderm areal density, *m* = *m_u_* + *m_b_* (Figure 4E). In the control, positive and negative velocity gradients on either side of the velocity peak (Figure 4D) cause mesoderm to separate and accumulate respectively as they extend towards the colony interior, predicting the emergence of a second density peak at later times (Figure 4E), primarily comprising basal mesoderm (Figure S4G). Equal responses (*α* = 1) also produce a second peak, but overpopulate it (Figure S4C). In the knockdown, undirected motility alone reproduces the evolution of the single-peaked areal-density profile in shAPLNR gastruloids (Figure 4E).

The model also captures the similar overall migration speeds of control and shAPLNR mesoderm (Figure 3E). The model reproduces spatial progression of radial speed (the magnitude of radial cell movement) observed in control gastruloids but does not capture its increase over time because it assumes a constant effective diffusivity. With 10 minutes between frames, undirected motility alone predicts an apparent radial speed, *|*Δ*r|/*Δ*t*, of *≈* 15*µ*m*/*h (Figure 4F), exceeding peak directed motility (Figure 4D). While uniform undirected motility dominates radial speed in both conditions, the directed motility from the proposed guidance mechanism is quantitatively sufficient to reproduce the APLNR-dependent mesoderm dispersal.

### APLNR depletion preserves broad paraxial and lateral mesoderm differentiation competence

To test whether APLNR is required for mesoderm specification independently of spatial migration, we used established directed differentiation protocols along two trajectories: anterior PS to paraxial mesoderm (APS–PM) and mid PS to lateral mesoderm (MPS–LM)^42^. Cultures were collected at days 1 and 2 (Figure 5A). These protocols use distinct combinations of WNT, FGF, Activin/NODAL, and BMP signaling to recapitulate signaling differences that pattern the PS along the anterior–posterior axis in vivo and subsequently generate paraxial or lateral mesoderm. For example, MPS induction includes a higher WNT dose at day 1, consistent with the posteriorizing role of WNT.

**Figure 5.**
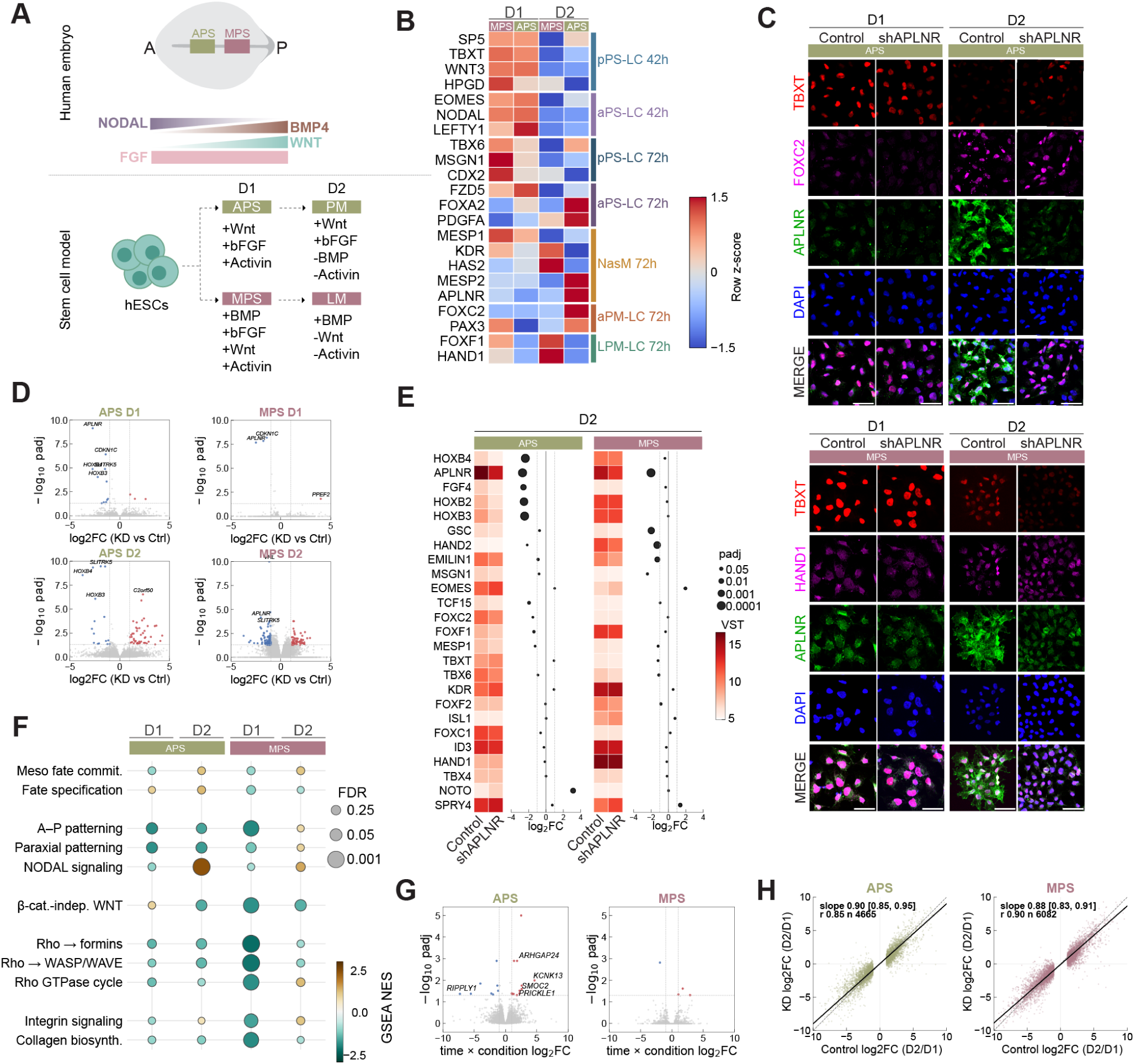
APLNR depletion preserves broad paraxial and lateral mesoderm differentiation competence. **(A)** Schematic of the directed-differentiation design. The doxycycline-inducible shAPLNR tNeon hESC line was differentiated toward paraxial mesoderm through an anterior PS intermediate (APS–PM) or lateral mesoderm through a mid PS intermediate (MPS–LM). *N* = 3 independent differentiations per arm, time point, and condition. **(B)** Expression of 20 genes associated with the indicated gastruloid states in control APS–PM and MPS–LM differentiations. Columns, mean across *N* = 3 control differentiations at APS D1, MPS D1, APS D2, and MPS D2; values are gene-wise *z*-scores across the four conditions; genes grouped by gastruloid state. **(C)** Representative immunofluorescence of APS–PM (top) and MPS–LM (bottom) cultures at days 1 and 2: APS–PM stained for TBXT, FOXC2, APLNR, and DAPI; MPS–LM for TBXT, HAND1, APLNR, and DAPI. Scale bars, 50 *µ*m. *N* = 3 independent differentiations. **(D)** Volcano plots of the shAPLNR-versus-control contrast at APS D1, MPS D1, APS D2, and MPS D2. Red and blue points, adjusted *p <* 0.05 and *|* log_2_ FC*| ≥* 1 (15, 10, 67, and 159 genes, respectively); selected genes labeled. Full results in Table S4. *N* = 3 independent differentiations. **(E)** Selected mesoderm fate, patterning, and signaling genes at day 2 during APS–PM and MPS–LM differentiation. Heatmap color, mean VST expression in control and shAPLNR cultures; adjacent plots, shAPLNR-versus-control log_2_ fold change, with dot size indicating adjusted *P* value. *N* = 3 independent differentiations. **(F)** Preranked gene-set enrichment of the shAPLNR-versus-control effect at APS D1, APS D2, MPS D1, and MPS D2, for selected Gene Ontology Biological Process gene sets grouped by category. Dot color, normalized enrichment score; dot size, false-discovery rate. Full results in Table S5. *N* = 3 independent differentiations. **(G)** Volcano plots of the time-by-condition interaction for APS–PM and MPS–LM differentiation (whether the effect of shAPLNR differs between day 1 and day 2). Colored points, adjusted *P <* 0.05 and *|*interaction log_2_ FC*| ≥* 1; selected genes labeled; genes with baseMean *<* 15 excluded from visualization. *N* = 3 independent differentiations per arm, time point, and condition. **(H)** Day-1-to-day-2 transcriptional changes in control and shAPLNR cultures for APS–PM and MPS–LM differentiation, for genes changing significantly between day 1 and day 2 in control (*p*_adj_ *<* 0.05, *|* log_2_ FC*| ≥* 1). Points, genes; dashed lines, *y* = *x*; solid lines, Deming regression fits, with slopes and 95% bootstrap confidence intervals within each panel (Methods). APS–PM, *n* = 4,665 genes; MPS–LM, *n* = 6,082 genes. *N* = 3 independent differentiations per arm, time point, and condition.

Comparison with gastruloid transcriptional states show that the directed differentiations recapitulates the expected PS and mesodermal programs (Figure 5B). At day 1, PS and nascent mesoderm-associated programs are broadly shared across both trajectories, consistent with a common early mesodermal state. By day 2, lineage-specific differences become more apparent, with APS–PM cultures enriched for paraxial mesoderm-associated genes such as *FOXC2* and MPS–LM cultures enriched for lateral mesoderm-associated genes such as *FOXF1*.

Control and shAPLNR cultures follow the expected APS–PM and MPS–LM differentiation trajectories (Figure 5C). Transcriptomes cluster primarily by time point and differentiation arm rather than knockdown condition (Figure S5A–C). Relatively few genes are differentially expressed at day 1, although *HOXB3* and *HOXB4* are significantly downregulated in APS cultures. Differential expression expands by day 2 to genes associated with developmental patterning, signaling, and morphogenesis (Figure 5D). APLNR depletion reduces expression of the anterior–posterior patterning genes *HOXB2*, *HOXB3*, and *HOXB4* in APS–PM cultures^43,44^. *FGF4* is also down-regulated whereas the axial mesoderm-associated gene *NOTO* is upregulated. MPS–LM cultures show reduced expression of *GSC*, *HAND2*, and *EMILIN1*. Core mesoderm and lineage-associated genes such as *TBXT*, *FOXC2*, and *FOXF1* nevertheless remain expressed along their expected differentiation trajectories (Figure 5E; Figure S5E).

Gene set enrichment analysis and inspection of leading-edge genes identified transcriptional programs affected by APLNR depletion (Figure 5F; Table S5). Mesoderm commitment and specification gene sets are not negatively enriched. Pathways associated with anterior–posterior and paraxial mesoderm patterning, non-canonical WNT and RHO signaling, integrin signaling, and extracellular-matrix organization instead show condition- and time-dependent changes.

APLNR depletion could also alter the progression of differentiation over time. Both control and shAPLNR cultures show extensive transcriptional changes from day 1 to day 2 along each differentiation trajectory (Figure S5D). A formal time-by-condition analysis identified a limited set of genes whose temporal responses differ in shAPLNR cells, with most changes occurring in APS–PM differentiation (Figure 5G). These included genes associated with developmental patterning and cell behavior, such as *RIPPLY1*, *PRICKLE1*, *ARHGAP24*, *SMOC2*, and *KCNK13*. Consistent with this, the overall day one to two transcriptional changes are highly similar between control and shAPLNR cultures in both differentiation arms (Figure 5H). Although the magnitude of some responses were modestly reduced following APLNR depletion, the overall direction and structure of the differentiation programs are preserved. Together, these analyses indicate that APLNR depletion selectively alters patterning, signaling, and morphogenetic programs while preserving overall paraxial and lateral mesodermal competence.

### APLNR loss alters mesoderm patterning and spatial progression in gastruloids

Having established that APLNR-depleted cells retain mesodermal competence under directed differentiation conditions, we examined mesoderm patterning in control and shAPLNR gastruloids by single-cell RNA sequencing at 42 and 72 h. The major transcriptional states are reproducibly recovered across biological experiments (Figure S6A–C). Control and shAPLNR gastruloids contain the same major PS-like and nascent mesoderm states, although their relative proportions differ (Figure 6A).

**Figure 6.**
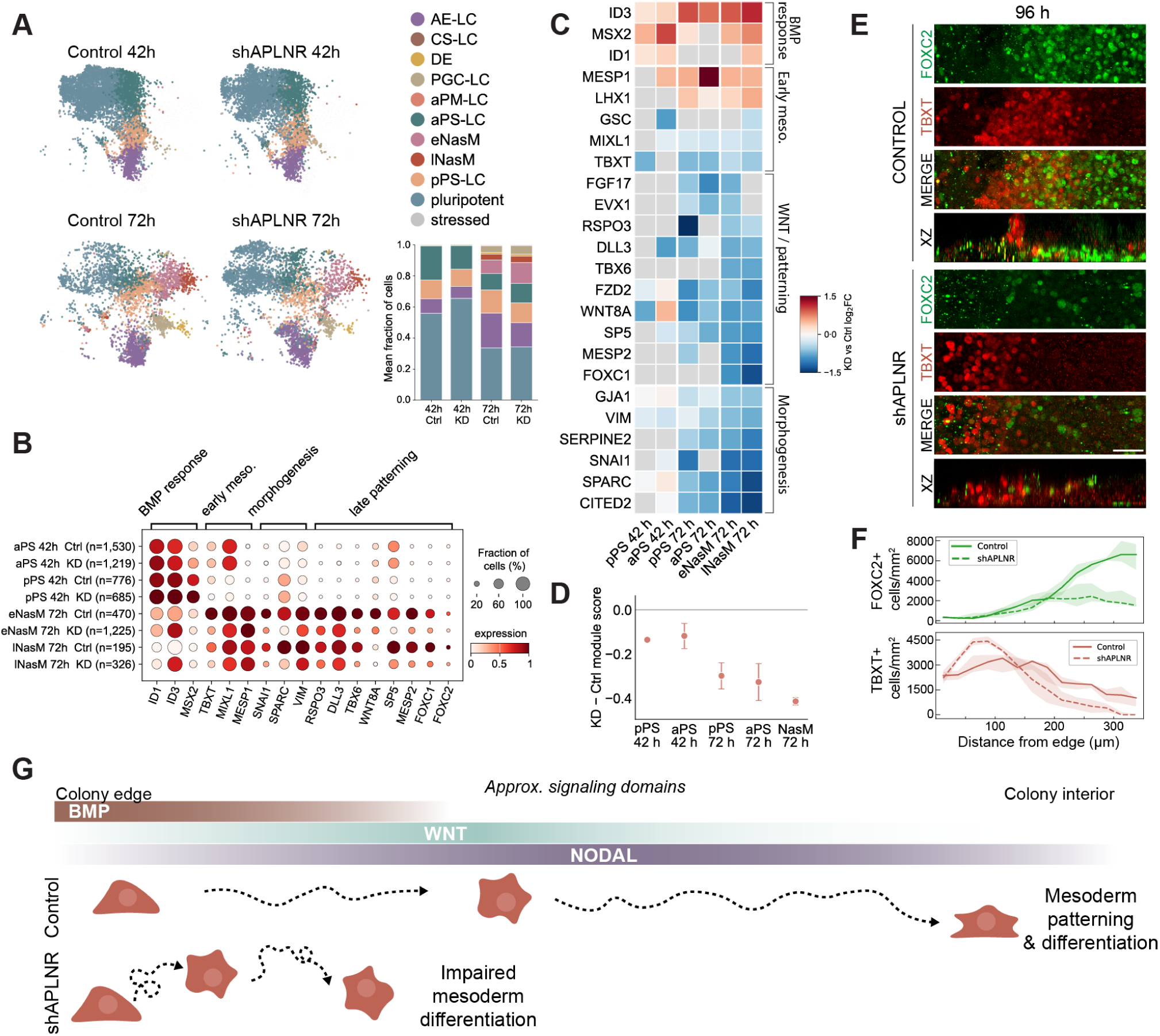
APLNR depletion alters mesoderm patterning and central paraxial organization in gastruloids. **(A)** UMAP representation of single-cell RNA-sequencing data from control and shAPLNR gastruloids at 42 and 72 h, colored by annotated cell state. Stacked bars show the mean fraction of cells assigned to each state by condition and time point. AE-LC, amniotic ectoderm-like; aPS-LC, anterior PS-like; pPS-LC, posterior PS-like; eNasM, early nascent mesoderm; lNasM, late nascent mesoderm; DE, definitive endoderm; PGC-LC, primordial germ cell-like. *N* = 2. Cell numbers: control 42 h, 6,746; shAPLNR 42 h, 5,487; control 72 h, 3,863; shAPLNR 72 h, 3,863 cells subsampled from 13,323. **(B)** Dot plot of selected BMP-response, early-mesoderm, morphogenesis, and late-patterning genes in aPS-LC and pPS-LC at 42 h and eNasM and lNasM at 72 h. Dot size, fraction of cells expressing the gene; color, scaled expression. Cell numbers per state and condition on the y-axis. *N* = 2. **(C)** Pseudobulk differential-expression heatmap of log_2_ fold change in shAPLNR relative to control within matched PS-like and nascent-mesoderm states at 42 and 72 h; genes grouped by BMP response, early mesoderm, WNT/patterning, and morphogenesis-associated programs. Full results in Table S6. *N* = 2. **(D)** Difference in the APLNR-containing WGCNA module score (defined in Figure 2G; *APLNR* excluded) between shAPLNR and control within the indicated PS-like and nascent-mesoderm populations. Points, shAPLNR–control difference; whiskers, variation across biological repeats. *N* = 2. **(E)** Representative immunofluorescence of FOXC2 and TBXT in control and shAPLNR gastruloids at 96 h; merged images below individual channels. Scale bar, 50 *µ*m. *N* = 1 independent experiment. **(F)** Radial density of FOXC2^+^ and TBXT^+^ segmented objects at 96 h in control and shAPLNR gastruloids, in 25-*µ*m annuli by distance from the colony edge (0 *µ*m, edge; 350 *µ*m, center), normalized to annular area. Lines, median; shading, IQR. Control, *n* = 5; shAPLNR, *n* = 4; *N* = 1. **(G)** Proposed model linking APLNR-dependent mesoderm migration to subsequent patterning. Approximate BMP, WNT, and NODAL signaling domains are shown schematically based on published work^25,46^; band extents do not represent measurements in this study.

shAPLNR cells retain expression of early mesoderm markers including *TBXT*, *MIXL1*, and *MESP1* but differ from controls in BMP-response, mesoderm patterning, and morphogenesis-associated genes (Figure 6B). Pseudobulk analysis across biological replicates shows increased expression of BMP-response genes including *ID3* and *MSX2* at 42 h. By 72 h, shAPLNR gastruloids show reduced expression of paraxial/presomitic mesoderm patterning genes including *TBX6*, *MESP2*, and *FOXC1*, alongside morphogenesis-associated genes such as *VIM*, *SNAI1*, and *SPARC* (Figure 6C). Neither *TBX6* nor *FOXC1* is significantly altered by APLNR depletion during APS–PM or MPS–LM directed differentiation. Interestingly, the organizer-associated transcription factor *GSC* ^45^ is downregulated both in lNasM at 72 h and during MPS–LM directed differentiation (Figure 5E; Figure 6C). This shared response links APLNR depletion to altered expression of a developmental patterning regulator in both systems.

The early increase in BMP-response genes is consistent with retention of shAPLNR mesoderm within the peripheral PS-like region (Figure 3H). BMP activity is highest near the gastruloid edge^24^, and previous measurements show that pSMAD1 activity in TBX6^+^ mesoderm decreases with increasing distance from the edge^46^. Reduced inward dispersal may therefore prolong mesoderm exposure to BMP signaling.

To examine coordinated changes in mesoderm developmental programs, we scored cells using the APLNR-containing co-expression module, excluding *APLNR* itself (Figure 2G; Table S3). Module scores are reduced in shAPLNR PS-like and nascent mesoderm populations, including nascent mesoderm at 72 h, consistent with coordinated attenuation of mesoderm developmental programs (Figure 6D).

The changes in paraxial mesoderm patterning genes at 72 h prompted us to examine later paraxial organization. Prior work identified FOXC2^+^ paraxial mesoderm beneath the colony interior during extended micropattern differentiation^30^. At 96 h, control gastruloids contain abundant FOXC2^+^ cells in the colony interior, whereas shAPLNR gastruloids show reduced central FOXC2^+^ enrichment despite retaining abundant TBXT^+^ cells (Figure 6E; Figure S6D). Radial cell-density profiles quantify this reduction in central FOXC2^+^ enrichment and show that TBXT^+^ cells remain concentrated in the PS-like region in shAPLNR gastruloids (Figure 6F). Together, these results support a model in which APLNR-dependent migration allows mesoderm to leave the peripheral PS-like region and encounter distinct signaling environments that shape subsequent mesoderm patterning (Figure 6G).

## DISCUSSION

We identify APLNR as a regulator of directed human mesoderm migration and nominate spatially organized APELA as a candidate guidance cue. During gastrulation, mesoderm progenitors must become motile while also interpreting spatial information that organizes their movement away from the PS. In endothelial cells, the APLNR ligand APLN promotes filopodia formation and vascular sprouting, whereas disruption of APLNR signaling reduces endothelial protrusions and impairs sprout progression^47^. During zebrafish gastrulation, the alternative APLNR ligand APELA/Toddler promotes mesendoderm motility and subsequently guides directed migration through a self-generated ligand gradient^10,12^. Our findings support a conserved role for APLNR in directed migration. This differs somewhat from zebrafish, where APELA/Toddler also acts as a motogen that increases random motility in addition to promoting directional migration^12^. In human gastruloids, APLNR loss disrupted inward directional bias while migration speed was only modestly reduced. This small speed change did not account for the dispersal defect, suggesting that APLNR primarily contributes to guidance in this context. Moreover, the complementary organization of APELA-expressing epithelial cells and APLNR expressing mesoderm provides a plausible spatial mechanism for orienting this response. APELA also exhibits approximately 25-fold greater potency than Apelin-13 in a recently developed APLNR conformation biosensor assay, further supporting its candidacy as a guidance ligand^48^. A minimal model of guidance up an APELA gradient was sufficient to reproduce the observed migration dynamics, whereas incorporating a self-generated gradient^12,49,50^ added parameters without being required to explain the data. Whether endogenous APELA forms a spatial gradient within human gastruloids remains an important question.

Our findings distinguish the requirement for APLNR in mesoderm guidance from the intrinsic capacity of progenitors to differentiate. Previous animal studies showed that disruption of APELA–APLNR signaling reduces mesodermal and cardiovascular derivatives, establishing an important developmental role for the pathway without resolving whether these phenotypes arise from defective migration or direct regulation of lineage specification^20,51^. Studies in human pluripotent stem cells have also implicated APLNR in mesoderm commitment and migration, as well as subsequent cardiomyocyte differentiation and contractile function^14,52^. Here, progenitors lacking APLNR retained broad competence to acquire paraxial and lateral mesoderm identities when exposed to direct instructive signals. The consequences of APLNR loss are therefore context dependent: lineage competence is largely preserved under uniformly supplied differentiation cues, whereas mesoderm organization is disrupted within a spatially patterned tissue. During early mesoderm development, APLNR is required for directed migration but not for initial mesoderm specification itself. This guidance may allow competent progenitors to move into distinct signaling environments. Future work will be required to determine how APLNR dependent migration and its roles in later lineage development are related.

Our study identifies APLNR-dependent directed migration as a contributor to human mesoderm patterning. These findings support a model in which migration allows mesoderm progenitors to leave the PS-like region and encounter distinct signaling environments as differentiation proceeds. In this model, movement links the spatial organization of signals to the developmental programs of responding cells. Directed migration therefore contributes to mesoderm development beyond redistributing cells: it helps position competent progenitors within environments that shape their subsequent patterning.

### Limitations of the study

Several limitations define the interpretation of this work. Although layer resolved imaging and orthogonal sections support basal dispersal of TBXT^+^ cells, limited z-sampling prevented continuous tracking of individual cells during their transition from the upper to the basal compartment. Expression data support APELA as a candidate guidance ligand, its endogenous spatial distribution and the respective contributions of APELA and Apelin remain unresolved. Direct visualization of active ligand gradients is challenging because ligand degradation can reduce bioactivity without a corresponding loss of fluorescence^53^. Our guidance model shows that a plausible ligand field can reproduce the observed dynamics but cannot distinguish between source derived and self-generated gradients or other modes of spatial APLNR activation. Recently developed APLNR conformation biosensors combined with ligand-specific perturbations could help resolve the guidance mechanism in intact gastruloids^48^.

Finally, altered signaling exposure following APLNR loss is inferred from mesoderm retention near the PS-like region and associated transcriptional changes rather than measured directly in migrating cells. Because BMP, WNT, and NODAL form a coupled signaling network in micropatterned gastruloids^25,27,28^, elevated BMP response transcription cannot be attributed to BMP signaling alone as the cause of later patterning defects. Simultaneous measurements of migration and signaling activity will be needed to establish this relationship. Furthermore, the radial organization of gastruloids does not fully reproduce the geometry or extraembryonic signaling environment of the human PS, limiting direct correspondence with mesoderm migration in vivo.

## RESOURCE AVAILABILITY

### Lead contact

Requests for further information and resources should be directed to and will be fulfilled by the lead contact, Margaret Gardel.

### Materials availability

This study did not generate new plasmids for public deposition; the shAPLNR-expressing cell line generated in this study is available from the lead contact upon request. Cell lines obtained from external sources — the Allen Institute for Cell Science SOX2–GFP and ZO-1–GFP reporter lines (via Coriell) and the TBXT–mNeonGreen H9 reporter line (gift of Valerie M. Weaver) — are available from their original sources as cited in the Key Resources Table.

### Data and code availability

- Bulk RNA-seq data have been deposited at GEO: GSE349990 and are publicly available as of the date of publication. Single-cell RNA-seq data have been deposited at GEO: GSE350897 and are publicly available as of the date of publication.
- All original code, including sequencing and chemotaxis-model analysis, has been deposited at https://github.com/amanda-brooke/aplnr_code and is publicly available as of the date of publication.
- Any additional information required to reanalyze the data reported in this paper is available from the lead contact upon request.

## ACKNOWLEDGMENTS

A.B.J. was supported by an NIH/NIGMS Ruth L. Kirschstein Predoctoral Individual National Research Service Award (F31 GM157880-02) and by an NIH T32 Developmental Biology Training Grant (T32HD055164). S.S. acknowledges support from Schmidt Sciences LLC and the National Research Foundation Singapore under its NRF Fellowship (NRF-NRFF17-2025-0004). M.L.G. acknowledges support from NIH R35GM156484 and the National Science Foundation through the Physics Frontier Center for Living Systems (NSF PHY-2317138) and is a Biohub Investigator. M.S. acknowledges support from NSF PHY-2413073, NIH R35GM156889, and HFSP Early Career Grant RGEC31/2024. This research was made possible through the Allen Cell Collection, available from Coriell Institute for Medical Research.

## AUTHOR CONTRIBUTIONS

Conceptualization, A.B.J., S.S., A.P., M.S., I.H., and M.L.G.; methodology, A.B.J., S.S., L.Y., A.P., Z.Y., M.S., and I.H.; software, A.B.J., A.P., and Z.Y.; formal analysis, A.B.J. and A.P.; investigation, A.B.J. and L.Y.; writing–original draft, A.B.J., A.P., and M.L.G.; writing–review & editing, all authors; visualization, A.B.J., A.P., and Z.Y.; funding acquisition, M.L.G.; supervision, S.S., M.S., I.H., and M.L.G.

## DECLARATION OF GENERATIVE AI AND AI-ASSISTED TECHNOLOGIES IN THE WRITING PROCESS

During the preparation of this work, the authors used Claude (Anthropic) and ChatGPT (OpenAI) to improve the language and readability of author written text. The authors reviewed and edited all output and take full responsibility for the content of the publication.

## SUPPLEMENTAL INFORMATION INDEX

Videos S1–S7 and their legends

Figures S1–S6 and their legends in a PDF

Table S1: LIANA ligand–receptor ranking (CS7, CS8) and full marker gene lists by cell state and biological repeat

Table S2: Per-repeat cell-state proportions (control vs. shAPLNR, 72 h)

Table S3: APLNR-containing WGCNA module (218 genes), module membership (*kME*) and hub-gene rank

Table S4: Bulk RNA-seq differentially expressed genes, APS–PM and MPS–LM contrasts

Table S5: Gene-set enrichment analysis results, APS–PM and MPS–LM

Table S6: scRNA-seq pseudobulk differential expression, control vs. shAPLNR

Table S7: Single-cell RNA-seq quality-control metrics per sample

## SUPPLEMENTAL VIDEO TITLES AND LEGENDS

**Video S1 (related to Figure 1C)**

Live imaging of a ZO-1–GFP reporter gastruloid from 0 to 48 h post-BMP4, showing progressive remodeling of the junctional network from the colony edge inward.

**Video S2 (related to Figure 1D)**

Live imaging of a parental TBXT–mNeonGreen reporter gastruloid from 24 to 48 h post-BMP4, showing single-cell trajectories during PS-like-region establishment and mesoderm dispersal.

**Video S3 (related to Figure 1D)**

Live imaging of a SOX2–GFP reporter gastruloid from 24 to 48 h post-BMP4, showing single-cell trajectories within the epiblast-like compartment.

**Video S4 (related to Figure 1H,I)**

Z-stack live imaging of a TBXT–mNeonGreen reporter gastruloid, showing basal, middle, and apical layers to illustrate mesoderm ingression beneath the epiblast-like layer.

**Video S5 (related to Figure 3C)**

Live imaging of control and shAPLNR TBXT–mNeonGreen gastruloids from 24 to 48 h post-BMP4, showing reduced inward dispersal following APLNR knockdown.

**Video S6 (related to Figure 3I,J)**

Live imaging of control TBXT–mNeonGreen gastruloids from 24 to 48 h post-BMP4 showing basal and upper planes side by side.

**Video S7 (related to Figure 3I,J)**

Live imaging of shAPLNR TBXT–mNeonGreen gastruloids from 24 to 48 h post-BMP4 showing basal and upper planes side by side.

## STAR METHODS

All experiments were conducted using established human pluripotent stem cell lines previously approved for research use. No new human embryos, fetal tissue, or patient samples were generated for this study. All work was conducted in accordance with institutional guidelines for human pluripotent stem cell research.

### Experimental model and study participant details

#### Human pluripotent stem cell lines

The cell lines used in this study were the H9 human embryonic stem cell TBXT–mNeonGreen reporter line (gift of Valerie M. Weaver, University of California, San Francisco) and its doxycycline-inducible shAPLNR derivative (generated in this study), the SOX2–GFP (AICS-0074-026) and ZO-1–GFP (AICS-0023) human induced pluripotent stem cell reporter lines, developed at the Allen Institute for Cell Science (allencell.org/cell-catalog) and available through Coriell^31^, and the SCTi003-A human iPSC line (STEMCELL Technologies), used for representative immunofluorescence imaging (Figure 1A,B).

### Method details

#### Micropatterning

PDMS stamps were fabricated as described in^54^ and coated with 5% laminin to generate 500– 600 µm diameter circular patterns on glass-bottom dishes. For additional live imaging, CYTOO 96-well micropatterned plates (700 µm) were used following manufacturer instructions.

#### Gastruloid differentiation

Gastruloids were generated as described in^21^. Briefly, cells were dissociated with ReLeSR, plated onto 5% laminin-coated patterns in mTeSR+ + 10 µM Y-27632 for 10 min (PDMS) or 30 min (CYTOO), rinsed, and allowed to epithelialize for 24 h. Differentiation was initiated by adding 50 ng/mL BMP4. Time 0 corresponds to BMP4 addition.

#### DOX-inducible APLNR knockdown line

A TRE-based shAPLNR construct was integrated into TBXT–mNeonGreen hESCs using pXAT2 Cas9 and puromycin selection, following^39^. Cells were electroporated using the Neon system. Knockdown was validated by qPCR. Doxycycline was added at 1 µg/mL for at least 5 days before differentiation and maintained throughout differentiation; matched controls were cultured without doxycycline.

#### Mesoderm progenitor generation and APELA stimulation

For the VASP/F-actin stimulation assay (Figure 3K–M), control and shAPLNR mesoderm progenitor cells (TBXT–mNeonGreen–shAPLNR line, *±* doxycycline) were generated using a monolayer mesoderm-induction protocol. Cells were plated at day 0 in mTeSR with 10 µM Y-27632, and mesoderm induction was initiated at day 1 by switching to STEMdiff Mesoderm Induction Medium (MIM; STEMCELL Technologies, Cat# 05221). Day 1 corresponds to 24 h and day 2 to 48 h after plating; cells were stimulated with vehicle or 100 ng/mL APELA for 5 min at day 2 (48 h), then fixed and stained as described in Immunofluorescence, below.

Cell masks were generated from F-actin images and used to define a cortical region spanning 0.8–3.8 *µ*m inward from the cell boundary. Peripheral VASP intensity was quantified as the mean background-subtracted VASP fluorescence within this cortical region. Cortical F-actin enrichment was calculated for each cell as the mean F-actin intensity within the cortical region divided by the mean F-actin intensity across the whole-cell region. Cell aspect ratio was calculated from the actin-derived cell mask. For comparisons of APELA-induced responses across independent experiments, Δ cortical F-actin enrichment and Δ aspect ratio were calculated for each APELA-treated cell relative to the median vehicle value from the same experiment and genotype.

#### Quantitative PCR

RNA (all samples *≥*100 ng/µL) was extracted with the Nucleospin RNA kit and reverse-transcribed with SuperScript IV VILO Master Mix. Resulting cDNA was diluted to 50 ng/µL in a 200 µL working stock, with 1 µL used as template per well. APLNR transcript was quantified by TaqMan qPCR (Applied Biosystems QuantStudio; TaqMan Fast Advanced Master Mix; APLNR assay Hs00270873 s1) with 18S (Hs99999901 s1) as the reference gene, and expression was calculated relative to matched dox-untreated control by the ΔΔ*C_t_* method, in *N* = 4 biological replicates.

#### Live imaging

Live imaging was performed on a Zeiss LSM 980 Airyscan using 8Y mode and a 20*×*/0.8 NA objective, with a stage-top incubator maintained at 37°C and 5% CO_2_. Images were acquired at 10-min intervals with 3.5-µm spacing between Z-planes.

#### Immunofluorescence

Samples were fixed in 4% PFA (15 min), permeabilized in 0.2% Triton X-100, blocked in 5% BSA, and incubated with primary antibodies overnight at 4°C (catalog numbers listed in Key Resources Table). Alexa Fluor-conjugated secondary antibodies were applied for 1 h at room temperature. Imaging was performed on the same Zeiss LSM 980 Airyscan system described above.

#### ZO-1 tight-junction quantification

Live ZO-1–GFP gastruloids were imaged for 48 h post-BMP4 (10 min/frame) and analyzed as maximum-intensity projections. For each frame, the colony outline was segmented to define its center and radius, and skeletonized junctions were assigned a radial distance from the colony edge, calibrated to the 700 µm micropattern (350 µm radius). The junction network was detected with a Sato ridge filter, hysteresis-thresholded and skeletonized; to measure continuous junctions and exclude internalized ZO-1 clusters, objects were retained only if thin (local width *≤∼*4 µm) and continuous (length *≥∼*23 µm). Junction density (skeleton length per unit area) was computed within radial bands and normalized to each band’s mean over the first three frames (“fraction of onset”), so values report change relative to the intact starting network. To avoid artifacts from tissue detachment, the colony was divided into twelve angular sectors and any sector whose tissue coverage fell below 70% was excluded at that timepoint. Data are the median across *n* = 6 gastruloids (two experiments) for the kymograph and mean *±* SEM for radial-band time courses.

#### Single-cell tracking and trajectory extraction

Cells were detected using StarDist^55^ in FIJI/TrackMate^56–58^ and linked using Kalman filter tracking with a maximum linking distance of 35 µm. Tracks containing fewer than five edges were excluded from all migration analyses, including Figure 1L. Tracks were split at division events, and mother and daughter trajectories were analyzed separately. Tracking data were exported for downstream analysis.

#### Radial velocity and speed–distance analysis

For each gastruloid, the colony center was estimated from the midpoint of the spatial bounding box in each frame. Colony diameter was estimated from the maximum chord length of a convex hull constructed from outer-rim cells (at or above the 90th percentile of radial distance) in the first frame. A cell’s distance from the colony edge was calculated as the estimated colony radius minus its distance from the center, with zero at the edge and increasing values toward the center. For consecutive positions separated by Δ*t* = 10 min, speed was calculated as displacement magnitude divided by Δ*t*. Radial velocity was calculated as the displacement projected onto the inward-pointing radial unit vector, divided by Δ*t*; positive values indicate inward movement. Speed and radial velocity were summarized at the track level before aggregation within gastruloids. Persistence was calculated as net start-to-end displacement divided by total path length, with values from zero to one.

Trajectory orientation was defined relative to the inward radial direction, with 0 indicating inward and *±π* outward movement. D<u>irectional con</u>centration was summarized per gastruloid using mean resultant length, 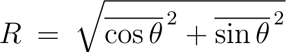. Rose plots show net start-to-end trajectory orientations in 15° bins. For Figure 1L, per-track mean speed was correlated with inward trajectory orientation within each gastruloid.

#### Basal and upper mesoderm compartments

Cells were assigned to layers according to nuclear-center Z-position in stacks acquired at 3.5-µm spacing. The lowest imaging plane was positioned at the nuclei nearest the glass substrate and defined the basal compartment; the remaining planes defined the upper compartment. Cells were tracked independently within compartments. Radial velocity was summarized per track within 2-h bins and then by the median within each gastruloid. Migration speed (Figure 3E) was summarized in 1-h bins; bins containing fewer than 10 gastruloids per condition were excluded.

#### Radial density profiles and PS-like boundary definition

To quantify the spatial extent of the TBXT^+^ cell population over time, tracked TBXT^+^ cells were binned by distance from the colony edge in 12.5 µm increments. Cell density per bin was computed and smoothed using a Gaussian filter (*σ* = 1 bin). Density profiles were computed separately for three time windows post-BMP4 addition: 24–36 h, 36–42 h, and 42–48 h.

A single, fixed PS-like boundary was defined from the 42–48 h density profile in control gastruloids. Because the dense PS-like region itself translocates inward over this period, defining the boundary from the final window ensures that cells classified as dispersed lie beyond the PS-like region at its most inward position, rather than reflecting displacement of the region as a whole. The PS-like peak was fit with a model that separates the Gaussian core from the inward-extending tail, and the boundary was set at *µ* + 1.5*σ* of the fitted core (*≈*127 µm from the colony edge; Figure S1B). This boundary was applied unchanged across all three time windows.

For the shAPLNR comparison (Figure 3H), the boundary was independently computed the same way from control gastruloids within that experiment (*µ* + 1.5*σ ≈* 112 µm) and applied to both control and shAPLNR gastruloids. As a robustness check, the shAPLNR reduction in dispersed fraction at 42–48 h, computed at the replicate level (*N* = 4), was confirmed using *µ* + 1*σ* (*≈*93 µm) and *µ* + 2*σ* (*≈*130 µm) as alternative thresholds. A track was classified as dispersed if it crossed inward beyond the threshold at least once during the specified time window. The dispersed fraction was the proportion of eligible tracks meeting this criterion within each gastruloid and time window. Comparisons across time windows used paired Wilcoxon signed-rank tests; control versus shAPLNR comparisons were analyzed as described in Quantification and statistical analysis. The dispersed front was defined by the 90th percentile of cells beyond the PS-like boundary, and bins with fewer than 10 dispersed cells were omitted.

#### PS-like region translocation

This section tracks the moving center of the PS-like region over time, distinct from the fixed dispersal boundary defined in Radial density profiles and PS-like boundary definition, above. For each 2-h time bin, the radial density profile of TBXT^+^ cells was computed as a function of distance from the colony edge (0–300 µm, 30 bins), with each track contributing a single value (its median radial position within the bin) to avoid weighting by track length. Because these per-bin profiles (unlike the single 42–48 h reference profile used for the boundary definition) include substantial numbers of already-dispersed cells at later times, each profile was fit by least-squares (scipy.optimize.curve fit) to a composite model consisting of a Gaussian component plus a one-sided inward exponential tail, rather than a Gaussian alone; the inward tail absorbs cells dispersing beyond the dense annulus so that the Gaussian center (*µ*) and width (*σ*) describe the PS-like region itself rather than being inflated by dispersed cells. The fitted center *µ* (in µm from the edge, with larger values corresponding to more interior/inward positions) was taken as the position of the PS-like region at each time point. For the trajectories shown in Figure 1I and Figure S3C, *µ* was computed per time bin and reported as the median across gastruloids, restricted to bins containing at least 10 gastruloids per group.

#### Minimal chemotaxis model

We model a radially symmetric, two-dimensional continuum on a disk of radius *R* = 350 *µ*m, reduced to solving equations in a one-dimensional radial coordinate *r ∈* [0*, R*]. We model dimensionless APELA concentration field *c*(*r, t*), rescaling the physical concentration by an unknown reference concentration so that the source has unit amplitude 1 h*^−^*^1^. We model mesoderm cell number densities *m_u_*(*r, t*) and *m_b_*(*r, t*) initially normalized to unit total mass over the two-dimensional domain. We let *r* = 0 define the colony center, such that distance from edge is *R − r*. Symmetry at the center requires *∂_r_c*(*r* = 0) = *∂_r_m_u_*(*r* = 0) = *∂_r_m_b_*(*r* = 0) = 0.

#### APELA concentration profile

We model ligand dynamics as

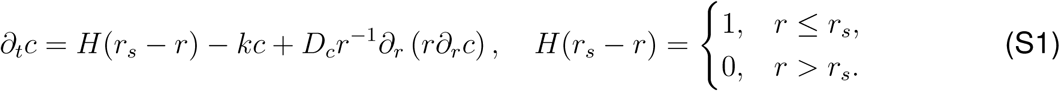

with a fixed ligand production rate for *r ∈* [0*, r_s_*], and uniform ligand clearance rate *k* and diffusivity *D_c_*. We impose an absorbing boundary condition *c*(*r* = *R*) = 0, representing leakage from the colony edge and surrounding medium^24,27^. This choice is motivated by evidence that leakage of a diffusible inhibitor from the colony edge contributes to edge fate patterning^21^. Our estimated ligand diffusivity (*D_c_* = 10^4^ *µ*m^2^*/*h) and clearance rate (*k* = 5 h*^−^*^1^) are comparable to estimates for rapidly diffusing NOGGIN in 2D gastruloids (*D* = 3.6 *×* 10^4^ *µ*m^2^*/*h, *k* = 3.6 h*^−^*^1^)^24^. Estimates for Toddler in zebrafish provide a similar order of magnitude reference (*D* = 1.8 *×* 10^5^ *µ*m^2^*/*h, *k* = 0.5 h*^−^*^1^)^12^. While a mesoderm-dependent uptake term might contribute to self-generated gradient dynamics^12^, boundary conditions and source geometry suffice to establish the gradient under a range of parameters (Figure S4A). We therefore model a uniform effective clearance rate for degradation and bulk leakage. Because this clearance rate is fast, and the onset of APELA production is uncertain, we model the steady state solution of Eq. (S1) rather than transient gradient formation.

#### Mesoderm migration model

We model upper and basal mesoderm, with densities *m_u_*(*r, t*) and *m_b_*(*r, t*), respectively:

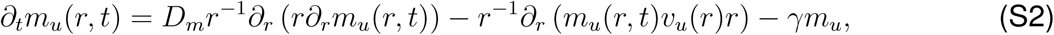

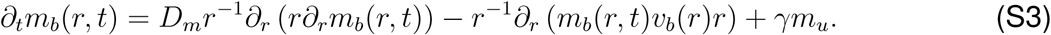

Here, *D_m_* is an effective diffusivity modeling undirected cell motility. We model directed motility as a mesoderm response to ligand concentration gradients with chemotactic velocity *v_b_*(*r*) = *β∂_r_c*(*r*) for basal mesoderm, and a weaker response *v_u_* = *αv_b_*, *α ∈* [0, 1] for upper mesoderm. The parameter *β* controls the strength of directed motility, and *α* the lesser response of the upper mesoderm. See Figure S4B for varying *D_m_* and *β*. Finally, the ingression parameter *γ* = 0.025 h*^−^*^1^ controls a fixed, irreversible transition rate between upper and basal populations, accounting for approximately half of the mesoderm cumulatively ingressing. See Figure S4C for varying *α* and *γ*. Total mesoderm areal density is initially distributed as,

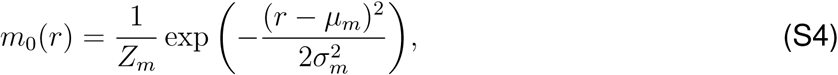

with normalization constant *Z_m_* such that 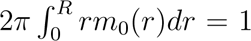 Initial condition parameters were rounded from Gaussian fits to the early 24 *−* 28 *h* density profiles of *N* = 4 control replicates (*µ_m_* = 320 *µ*m and *σ_m_* = 11 *µ*m) and shAPLNR replicates (*µ_m_* = 305 *µ*m and *σ_m_* = 12 *µ*m) (Figure S4F). All mesoderm is initially upper (*m_u_*(*r,* 0) = *m*_0_(*r*), *m_b_*(*r,* 0) = 0) and we impose a Dirichlet boundary condition *m_u_*(*R, t*) = *m_b_*(*R, t*) = 0, consistent with the low mesoderm density observed at the colony edge. Figure S4D shows similar dispersal behavior with a no-flux boundary condition. Because ligand concentration, receptor occupancy, and APELA/APLNR response parameters are not known in gastruloids, we used linear phenomenological response terms. A saturating response function could be required to model ligand overexpression, which can disrupt directed cell migration^59^. We neglect cell proliferation, as cell number increases plateau around the onset of directed migration (Figure 1E) and do not appear to impact the early mesoderm distribution (Figure 1F).

#### Model implementation

We solve Eqs. (S1)-(S3) on a uniform radial grid with *N_r_* = 200 points. The discretized steady-state of Eq. (S1) was solved directly. For mesoderm dynamics, diffusion was treated implicitly and chemotactic fluxes explicitly using an upwind discretization. During each timestep, a fraction 1 *− e^−γ^*^Δ^*^t^* of upper mesoderm is transferred to basal mesoderm. Simulations were run for *T* = 30 h, representing 24 h - 54 h after BMP4 addition. APLNR knockdowns were implemented by setting the chemotactic response parameter *β* = 0. Separate early-profile initial condition parameters were obtained for the APLNR knockdown, with *µ_m_* = 305 *µ*m and *σ_m_* = 15 *µ*m. All other dynamical parameters were shared.

#### Radial speed and velocity comparisons

Using the trajectories and radial coordinates described above, inward radial displacements Δ*r* were calculated by projecting each displacement across consecutive frames onto the inward-pointing radial unit vector at its midpoint. Radial speed was calculated as 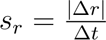. Speeds were averaged within 30 *×* 60 radial-position and time bins and combined across four gastruloids, excluding bins with fewer than five displacements. To compare speeds from discrete experimental trajectories with the continuum model, a model expected radial speed was computed accounting for both radial drift *v* = (*m_u_v_u_* +*m_b_v_b_*)*/*(*m_u_* +*m_b_*) and random motility 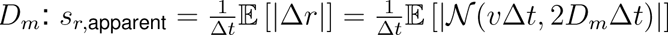, using the experimental frame interval Δ*t* = 10 minutes. Model bins below 5% of maximum density are masked. Similarly, inward radial velocities were calculated across consecutive frames as 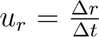 and in the model as: 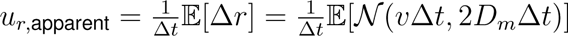. Means are computed only for frames and radial bins with at least two valid displacements. For each radial bin, temporal averages weight frames equally, and displayed bins require at least ten valid frames in each replicate.

#### Model sensitivity analyses

We assessed parameter sensitivity in the m<u>odel</u> by sweeping parameter pairs for ligand source radius *r_s_*and gradient length scale 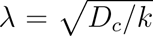 (varying *D_c_* with fixed *k*), chemotactic response *β* and mesoderm diffusivity *D_m_*, and upper response fraction *α* and ingression rate *γ* (Figure S4A–C). For each parameter combination, we calculated the normalized RMSE of mesoderm areal density across radial bins between model and experimental control density profiles averaged over 44–54 h. Baseline parameters lie in broad regions of low error, indicating that observed density dynamics do not require precise parameter tuning. Profiles from parameter combinations at the corners of each sweep illustrate alternative model behavior. Figure S4A shows that the source distribution, here modeled simply as a Heaviside function, is important for reshaping the initially symmetric mesoderm density profile, with best fits at an intermediate source radius and gradient lengthscale, and inward dispersal over a wide range of parameters. Figure S4B demonstrates that trading diffusivity for chemotactic response can preserve similar density profiles, so long as *β* is sufficiently high. Finally, ingression has the slowest modeled timescale, and Figure S4C shows i) that the density data are also compatible with no upper mesoderm directed response (*α* = 0) and ii) that a second mesoderm density peak can be generated without distinct behavior for upper and basal mesoderm (*α* = 1), but in this case the inner peak density exceeds the outer.

| Parameter | Name | Value |
| --- | --- | --- |
| $r_s$ | APELA source radius | 220 $\mu\text{m}$ |
| $k$ | APELA clearance rate | 5 $\text{h}^{-1}$ |
| $D_c$ | APELA diffusivity | 10<sup>4</sup> $\mu\text{m}^2/\text{h}$ |
| $D_m$ | Mesoderm diffusivity | 30 $\mu\text{m}^2/\text{h}$ |
| $\alpha$ | Upper-mesoderm response fraction | 0.5 |
| $\beta$ | Basal-mesoderm chemotactic response | 5 $\times 10^3$ $\mu\text{m}^2/\text{h}$ |
| $\gamma$ | Ingression rate | 0.025 $\text{h}^{-1}$ |

#### Directed differentiation toward paraxial and lateral mesoderm

Cells were differentiated through anterior primitive streak to paraxial mesoderm (APS–PM) or through mid primitive streak to lateral mesoderm (MPS–LM) using the protocols of^42^. Control and shAPLNR cultures were collected at days 1 and 2. Doxycycline-treated cultures received 1 µg/mL doxycycline for at least 5 days before induction and throughout differentiation. Three independent differentiations were performed per trajectory, time point, and condition.

### Bulk RNA-sequencing

#### Samples and design

Paired-end bulk RNA-seq was performed on hESC-derived anterior (APS) and mid (MPS) PS populations at two differentiation days (24 and 48 h), each with a doxycycline-inducible APLNR knockdown (shAPLNR) and a matched control, in *N* = 3 biological replicates per condition (R1–R3; 24 libraries).

#### Sequencing QC

Per-sample read quality was assessed with FastQC and aggregated with MultiQC by the sequencing core, including an rRNA-content check; for QC, reads were mapped to hg38. Genome mapping was used for QC only — transcript quantification used kallisto against the GENCODE transcriptome (below).

#### Quantification

Reads were pseudoaligned and transcript abundances estimated with kallisto v0.46.1 against the GENCODE v46 (GRCh38) transcriptome (gencode.v46.transcripts.fa. gz; 254,070 transcript targets); libraries were unstranded. Transcript-level estimates were summarized to genes with tximport v1.30.0 using the GENCODE v46 transcript-to-gene map.

#### Differential expression

Analysis used DESeq2^60^. Counts were imported with DESeqDataSetFromTximport under the design *∼* replicate + group, where group encodes the cell-type *×* day *×* condition combination, and genes were pre-filtered to those with *≥* 10 counts in *≥* 4 samples. The variance-stabilizing transform (vst, blind = FALSE) was used for visualization and PCA; for PCA, replicate batch was removed with limma removeBatchEffect under a *∼* celltype *∗* day *∗* condition model. Contrasts extracted from the fitted model were: the knockdown effect (shAPLNR vs control within each celltype *×* day), the temporal program (48 vs 24 h within each celltype *×* condition), the celltype comparison (APS vs MPS within each condition *×* day), and the corresponding interaction terms. Because replicate is included in the design, reported log2 fold-changes are DESeq2 model coefficients adjusted for replicate batch (unshrunken maximum-likelihood estimates) rather than ratios of normalized counts. Genes were called differentially expressed from Wald tests at *p*_adj_ *<* 0.05 (Benjamini–Hochberg) and *|*log2FC*| ≥* 1.

#### Gene-set enrichment

Preranked GSEA^61^ was performed with GSEApy (prerank), ranking genes by the DESeq2 Wald statistic, against MSigDB v2026.1 Hallmark, GO Biological Process (c5.go.bp) and Reactome (c2.cp) collections (gene symbols). Gene sets were restricted to 15–500 genes; 1000 permutations; seed 149; weighted (weight 1.0) enrichment. Terms were considered significant at FDR *<* 0.25; developmental, patterning, migration, adhesion and matrix terms were highlighted from the GO:BP results.

#### Figure-level analysis

Figures were generated in Python (numpy, pandas, matplotlib, scipy) from the DESeq2, expression and GSEA outputs. For symbol-level and unbiased analyses, duplicate gene symbols (multiple Ensembl IDs) were resolved by retaining the highest-expressed gene entry, and ribosomal-protein, mitochondrial and rRNA genes were excluded from unbiased variable-gene selection. For the trajectory analysis, among genes with a significant temporal program in control, the per-gene 48-vs-24 h log2 fold-change under shAPLNR was compared with control: concordance is reported as Pearson *r*, and the magnitude (slope) estimated by Deming regression (errors-in-both-variables), with the error-variance ratio set from the DESeq2 standard errors and a 95% CI from 500 gene-level bootstraps.

### Single-cell RNA sequencing

#### Sample preparation and sequencing

Two independent experiments were performed, one at the University of Chicago and one at the University of Michigan, using the same TBXT– mNeonGreen–shAPLNR cell line. Doxycycline-treated cells received doxycycline for at least 5 days before BMP4 induction and throughout differentiation, alongside matched untreated controls. Samples were collected at 42 and 72 h, with approximately 6,000 cells targeted per time point per experiment across control and doxycycline-treated samples. Samples were harvested using the 10x Flex fixation kit according to the manufacturer’s instructions. Single-cell RNA sequencing was performed by the University of Michigan Advanced Genomics Core. Fixed dissociated cells were counted using a Luna FX7 (Logos Biosystems). Probe hybridization, sample pooling, and library preparation followed the manufacturer’s instructions for Chromium Fixed RNA Profiling Reagent Kits for Multiplexed Samples (10x Genomics, PN 1000568). Library quality was assessed using a LabChip GXII HT (PerkinElmer), and libraries were quantified using Qubit (Thermo Fisher Scientific). Pooled libraries underwent paired-end sequencing on an Illumina NovaSeq X Plus. BCL Convert (Illumina) generated demultiplexed FASTQ files, and Cell Ranger 7.0.1 (10x Genomics) generated count matrices using the supplied human GRCh38 reference and default settings.

#### Quality control and normalization

Initial preprocessing retained cells with at least 500 detected genes, at least 1,000 UMIs, and no more than 20% mitochondrial counts. Cells exceeding the sample-specific median plus five median absolute deviations for detected genes or total counts were excluded. Final downstream cell selection used a saved barcode list generated by an additional sample-specific QC pipeline. This pipeline applied UMI bounds separately to each sample, removed genes detected in fewer than 10 cells per sample, and retained cells with mitochondrial fractions no greater than 20%. UMI ranges for control/shAPLNR samples were 15,000–35,000/10,000–35,000 (Chicago, 42 h), 4,500–20,000/5,000–20,000 (Chicago, 72 h), 7,000–25,000/8,000–25,000 (Michigan, 42 h), and 5,000–30,000/4,000–20,000 (Michigan, 72 h). Sample labels originally recorded as 48 h were corrected to the actual collection time of 42 h. Raw counts were retained for pseudobulk analysis; expression used for visualization was normalized to 10,000 counts per cell and log1p-transformed.

#### Single-cell embedding and cell-state annotation

Single-cell datasets from both biological repeat experiments were analyzed in a common processed object using Scanpy^62^. Query cells and the published gastruloid reference^30^ were jointly integrated using scVI^63^ with raw counts and sample-level batch labels. The model used 30 latent dimensions, two hidden layers of 128 units, and a negative-binomial likelihood, with up to 250 training epochs and early stopping. Reference cell-state labels were transferred to query cells using a distance-weighted 25-nearest-neighbor classifier in the joint latent space. UMAP was computed from this representation using 20 neighbors and a minimum distance of 0.3. For comparisons of cell-state abundance, proportions were calculated independently within each biological repeat and experimental condition rather than from pooled cells, preventing differences in sequencing depth or recovered cell number from disproportionately weighting either repeat. For APLNR and APELA expression visualization, MAGIC^64^ (magic-impute v3.0.0; knn=5, t=1, decay=1, n pca=100, random state=0) was applied to log-normalized expression of the top 2,000 highly variable genes (Scanpy, flavor=’seurat’) plus the genes of interest. The diffusion operator was fitted separately for each gastruloid site and once across the embryo dataset.

Cell-state proportions are reported as the mean of the two biological repeats, each weighted equally. Because control and shAPLNR populations differ substantially in size (5,522 vs. 20,641 cells at 72 h), proportions rather than raw counts are compared, and only states whose repeats agree in direction are reported as changed. Per-state counts and per-repeat proportions are given in Table S2. Cell-state proportion changes were summarized descriptively using direction concordance across the two independent experiments; cells were not treated as independent biological replicates for significance testing.

#### Weighted gene co-expression network analysis and module scoring

The primary WGCNA network was built from control mesodermal cells, aggregated into pseudocells to reduce sparsity: within each biological repeat *×* timepoint *×* cell-state stratum, cells were *K*-means clustered to *∼*25 cells per pseudocell (minimum 10), and raw counts were summed, converted to CP10K, and log1p-transformed.

A signed network (PyWGCNA; soft-thresholding power 14, signed topological overlap matrix) was built from the pseudocell matrix; modules were called by hierarchical clustering/dynamic tree cutting (minimum size 30 genes) and merged at an eigengene dissimilarity threshold of 0.20. The module containing *APLNR* — the receptor of interest — was carried forward: a 218-gene module with *APLNR kME* = 0.83.

As a robustness check, an independent network built from all control cell states (rather than mesoderm alone) yielded a separate 151-gene APLNR-containing module, sharing 81 genes with the primary module; *kME* values for these shared genes were strongly correlated between the two networks (*r ≈* 0.91; Figure S2G).

For KD cells, the network was not rerun; the frozen control-defined module (217 of 218 genes, excluding *APLNR* itself so depletion could not trivially lower the score) was applied to the single-cell matrix (duplicate symbols summed, CP10K, log1p) with each gene *z*-scored to its control mean/SD. The per-cell score was the mean *z*-score across the 217 genes, summarized by biological repeat, timepoint, cell state, and condition (eNasM/lNasM merged for the nascent mesoderm comparison); KD–control differences were calculated within each repeat and then summarized across repeats (Figure 6D).

Module preservation in shAPLNR cells was tested with WGCNA’s modulePreservation (control assignments as reference, shAPLNR pseudocells as test network; signed, Pearson, 200 permutations, seed 17; 3,000 genes, 101 pseudocells/condition): the module showed strong preservation (*Z*_summary_ = 25.921, median rank = 3), indicating its co-expression structure is retained, not disrupted, under knockdown.

#### Module gene-set enrichment

Preranked GSEA (GSEApy prerank) was performed on the APLNR-containing WGCNA module using signed module membership (*kME*) as the ranking statistic, against the Enrichr GO Biological Process 2023 and Reactome 2022 collections. The complete ranked list of genes retained in the mesoderm WGCNA analysis served as the gene universe; no separate background gene list was specified. Gene sets were restricted to 10–500 genes; analyses used 10,000 permutations. Positively enriched terms (normalized enrichment score *>* 0, FDR *q <* 0.05) were considered significant (Figure 2G; Figure S2H).

#### Gene–gene correlation

Pearson correlations of *APELA* and *APLNR* with other genes were calculated independently in each biological repeat on sequencing-depth-residualized normalized expression. The consensus correlation is the equal-weight Fisher *z*-transformed mean across repeats. Genes shown in Figure 2H are among the strongest associations with concordant direction in both repeats.

#### Marker dot plots

Marker expression was evaluated in the cell states, conditions, and time points displayed in each panel. Mean log1p-transformed, library-size-normalized expression and the fraction of cells with detectable expression were calculated from cells pooled across the two experiments within each displayed group. Dot color represents mean expression scaled from zero to one separately for each gene across the displayed groups (Scanpy, standard scale=’var’); dot size represents the fraction of cells with detectable expression.

#### State-resolved pseudobulk differential expression

Differential expression was evaluated separately within aPS-LC and pPS-LC at 42 h and eNasM and lNasM at 72 h. Raw single-cell counts were summed within each biological repeat *×* condition *×* cell-state combination to generate pseudobulk samples. State-resolved pseudobulk analyses were performed using Python and R. Control and APLNR-KD pseudobulks were compared using DESeq2, using a design that included biological repeat/site and condition to estimate the doxycycline-treated-versus-control effect within each state and time point. Genes were ranked according to the resulting KD-versus-control effect estimates, and selected positively and negatively affected genes were displayed in heatmaps.

### Spatial analysis of APLNR and APELA in Carnegie stage 7 and 8 human embryos

Published spatial transcriptomic datasets from Carnegie stage 7 (CS7) and Carnegie stage 8 (CS8) human embryos were obtained from^36,38^. For CS8, spatially registered Stereo-seq data comprised 38,562 spots across 62 serial sections. Anterior–posterior positions were taken from the deposited spatial registration and converted to approximate physical distance using the reported section spacing. Raw UMI counts for *APLNR* and *APELA* were extracted from the deposited count matrix and normalized to total transcriptomic library size per spot as

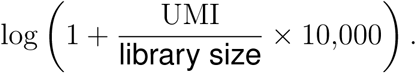

Tissue identities were based on the deposited CS8 annotations. Mesodermal populations were further classified using marker genes reported in the CS8 study, and related mesodermal subtypes, including axial, paraxial, lateral plate/intermediate, and extraembryonic mesoderm, were combined into a single mesoderm category for visualization. Anterior–posterior expression profiles were calculated as the mean normalized expression of each gene across embryonic spots within each serial section. Sections containing fewer than 20 embryonic spots were excluded, internal gaps were linearly interpolated between neighboring sections, and profiles were smoothed with a Gaussian kernel (*σ* = 2 sections).

CS7 spatial annotations and section coordinates were obtained from^38^. *APLNR* expression was observed directly; *APELA*, absent from the retained spatial gene set, was imputed from the Tyser et al. CS7 single-cell reference^35^. Datasets were projected into a joint PCA space using 1,483 shared genes. For each spatial observation, *APELA* expression was estimated as the inverse-distance-squared weighted mean of expression in its 15 nearest reference cells. Reconstruction of measured developmental genes (*APLNR*, *HAND1*, *BMP4*, and *TBXT*) yielded correlations of *r* = 0.71–0.80 with observed expression. Anterior–posterior profiles used the section-wise averaging and smoothing described above. Profiles were normalized independently by gene and stage and displayed on a common physical axis; absolute expression magnitudes were not compared between stages.

### APELA and APLN transcript contribution

Observed APELA and APLN counts were normalized to counts per million within each cell and summed by annotated state and time point in control gastruloids. States with fewer than 10 cells at either time point and stressed or ambiguous populations were excluded. Relative APLN contribution was calculated as APLN/(APELA + APLN) within each compartment. The same analysis was applied to the CS7 single-cell reference; CS8 spatial data were normalized per spot and aggregated by annotated compartment, excluding extraembryonic yolk-sac and visceral endoderm populations. These sums reflect both expression and sampled population size; absolute values were not compared across datasets.

### Quantification and statistical analysis

Analyses were performed using Python 3.11, NumPy, SciPy, pandas, statsmodels, and custom scripts, with R packages used for sequencing analyses as described above. *N* denotes independent experiments or biological replicates; *n* denotes gastruloids unless otherwise specified. Summary statistics and sample sizes are reported in the figure legends. Migration measurements were summarized per track and then per gastruloid. Comparisons of TBXT^+^ and SOX2^+^ migration speed used two-sided Mann–Whitney *U* tests. Comparisons across time windows within the same gastruloids used paired Wilcoxon signed-rank tests with Bonferroni correction for multiple pairwise comparisons. Per-gastruloid speed–orientation correlations were compared against zero using a one-sample Wilcoxon signed-rank test. Tests were two-sided unless otherwise specified, with *α* = 0.05.

For shAPLNR comparisons, gastruloids were nested within four independent biological replicates. Dispersed fraction (Figure 3H), defined by whether a track crossed the PS-like boundary at least once during the analysis interval, was analyzed using a linear mixed-effects model with condition as a fixed effect and a random intercept for biological replicate (*n* = 20 gastruloids per condition, *N* = 4). To assess whether reduced dispersal was explained by lower migration speed, the model was refit with each gastruloid’s mean migration speed as a covariate. Migration speed was analyzed using the same mixed-effects structure applied to per-gastruloid median speed. Replicate-level paired Wilcoxon signed-rank tests were also calculated for speed and dispersal as sensitivity analyses. Directional concentration and persistence were compared using replicate-level paired Wilcoxon signed-rank tests. Tracked TBXT^+^ cell numbers were compared using a paired *t*-test at the biological-replicate level. APLNR qPCR expression relative to control was compared against one using a one-sample *t*-test. PS-like region translocation and sequencing analyses were analyzed as described in their respective sections.

For shAPLNR imaging experiments, doxycycline-treated and untreated cultures were plated on the same plate and imaged simultaneously. Tracking and analysis settings were applied consistently across conditions. Biological replicate numbers varied by assay and are specified in the figure legends.

#### Use of AI-assisted tools

Large language models were used to assist with drafting and debugging analysis and plotting code. All analytical choices, statistical approaches, code validation, quantitative interpretation, and conclusions were reviewed and performed by the authors.

## Additional resources

Additional details, scripts, and reagents are available upon request.

